# Swiftly squeaky clean: lessons learned from eradicating an overpopulation of rats on an island of constraints

**DOI:** 10.1101/2025.02.19.639051

**Authors:** Tatiane Micheletti, Thayna J. Mello, Carlos Verona, Vinicius P. O. Gasparotto, Ricardo Krul, Ricardo Araujo, Thali Sampaio, Paulo Rogerio Mangini

**Affiliations:** Technische Universitat Dresden; Chico Mendes Institute for Biodiversity Conservation, ICMBio; Oswaldo Cruz Foundation (Fiocruz); Brazilian Institute for Conservation Medicine TRIADE

## Abstract

Invasive rats threaten island biodiversity, disrupting ecosystems and endangering native species. While rat eradication has succeeded on many islands, tropical islands present unique management challenges. Strict regulations and financial constraints on some tropical islands further limit proven eradication methods, complicating rodent management. This study addresses these challenges by demonstrating a real-time active adaptive management (RAM) approach, providing a cautious, cost-efficient, scientifically grounded, and adaptive pathway to eradication, while adhering to strict environmental regulations. We implemented RAM on a Brazilian island at USD 3,300 per hectare, integrating rodenticide (brodifacoum) application, population monitoring, and iterative management adjustments. This approach eradicated a rat overpopulation within five months, with early population increases observed in the threatened masked booby and the endemic Noronha elaenia and Noronha skink. Despite logistical constraints, RAM proved effective and cost-efficient, marking its first application in a biological system. Our findings highlight the value of innovation, interdisciplinary collaboration, and adaptive decision-making when the application of best-practice methods is constrained.

## 1 INTRODUCTION

Biodiversity conservation faces growing threats, from climate change to resource overexploitation (Rands et al. 2010). On islands, invasive species (IS) are a major challenge, vspreading rapidly and causing severe ecological disruptions (Blackburn 2004). Threatening local economies, public health, and cultural integrity (Nghiem et al. 2013), IS are closely linked to global biodiversity decline (Vitousek et al. 1997; Blackburn 2004). Among them, black rats (*Rattus rattus*) are particularly damaging due to their adaptability and high reproductive rates (Towns et al. 2006; Doherty et al. 2016). They prey on native species, compete for resources, and disrupt entire ecosystems, driving biodiversity loss (Towns et al. 2009; Fukasawa et al. 2013; Tabak et al. 2014; Harper & Bunbury 2015; Duron et al. 2017; Graham et al. 2018; Quiterie et al. 2019). However, rat eradication has proven feasible and highly effective for island conservation (Holmes et al. 2019).

Successful rat eradications have been documented worldwide, from Alaska (Kurle et al. 2021) to New Zealand (Russell & Broome 2016), though most occurred in subtropical (e.g., Howald et al. 2010) and temperate regions (e.g., Capizzi 2020). Despite notable exceptions (Samaniego-Herrera et al. 2018; Griffiths et al. 2019), black rat eradications on tropical islands are less common, with failure rates 2 to 2.4 times higher than in temperate zones (Russell & Holmes 2015). Ecologically, tropical conditions favor invasive species by providing year-round resources, mild winters (Holmes et al. 2015) , and reduced bait exposure due to competition with native fauna (Wegmann et al. 2011). Socioeconomic barriers, including high costs, limited expertise, weak collaboration, and regulatory constraints, further hinder success (Samaniego et al. 2021). While strict adherence to best practices improves outcomes (Samaniego et al. 2021), financial and regulatory limitations prevent their full implementation.

Overcoming these challenges requires innovative approaches that account for ecological, social, and economic complexities while maximizing eradication success. Adaptive management (AM) has emerged as a crucial strategy for improving conservation decisions (Chadès et al. 2017). Despite varying definitions (Westgate et al. 2013; Rist et al. 2013; Williams & Brown 2014; Månsson et al. 2023), its core principle remains integrating learning into management (McCarthy & Possingham 2007; McDonald-Madden et al. 2010; Chadès et al. 2017; Duron et al. 2017). Active AM follows an iterative six-step process (modified from Westgate et al. 2013): 1) defining clear management goals; 2) selecting multiple strategies; 3) measuring system responses; 4) implementing actions; 5) continuously monitoring outcomes; and 6) adjusting strategies accordingly. This cycle continuously incorporates new information, reducing uncertainties that could hinder eradication efforts.

With rapid ecosystem changes in recent decades, real-time monitoring has been increasingly integrated into AM, leading us to define ’real-time active adaptive management’ (RAM). While RAM has advanced in automated fields like water (Rao et al. 1992) and soil management (Park & Harmon 2011), and more recently epidemiology (Atkins et al. 2020), it remains underdeveloped in ecology and biological systems. Despite its recognized value, AM is rarely implemented in ecological projects, a trend persisting for over a decade (Westgate et al. 2013; Rist et al. 2013; Månsson et al. 2023).

We present a successful RAM project that rapidly eradicated a dense black rat population on a tropical island, overcoming ecological, financial, and regulatory challenges. To our knowledge, this is the first comprehensive application of AM (following Westgate et al. 2013) and the first to document real-time monitoring in a biological system. By detailing each step of the RAM framework and the challenges encountered, we provide a practical model for invasive species management where best practices are difficult to implement. This work highlights the value of interdisciplinary collaboration, adaptive decision-making, and cautious innovation, offering insights for future conservation and policy development.

## 2 METHODS

### 2.1 Study area and species

Ilha do Meio, part of the Fernando de Noronha (FN) archipelago (3°50’41’’S, 32°25’36’’W), is an 18-hectare uninhabited island of high ecological importance, free from human infrastructure (Appendix A, Figure A1). Comprising 21 islands and islets and located 345 km off Brazil’s northeastern coast, the volcanic archipelago is a Natural World Heritage site (UNESCO 2001) due to its importance as a breeding and feeding area for seabirds and endemic species. Ilha do Meio hosts the endangered insular land crab (*Johngarthia lagostoma*), as well as breeding populations of the masked (*Sula dactylatra*), brown (*Sula leucogaster*), and the endangered red-footed booby (*Sula sula*), along with the white-tailed tropicbird (*Phaethon lepturus*). The island also supports the endemic Noronha elaenia (*Elaenia ridleyana*), the endangered Noronha skink – locally known as mabuya (*Trachylepis atlantica*), and the Ridley’s worm lizard (*Amphisbaena ridleyi*). Neighboring Ilha Rata, a larger island 175 meters away separated by a strong tidal channel shares similar characteristics and was used as a control site for wildlife monitoring. While eradication was limited to Ilha do Meio, planned monitoring focused on the land crab, Noronha elaenia, mabuya, white-tailed tropicbird, and three booby species.

Fernando de Noronha has a warm climate, with water temperatures averaging 27°C and air temperatures ranging from 25°C to 31°C. Annual rainfall of 1,400 mm occurs mainly from January to July, with a drier season from August to December (WeatherSpark 2024). The vegetation is classified as Seasonal Deciduous Forest, varying between wet and dry seasons (Teixeira & Linsker 2003). Despite its protected status within the Marine National Park, the archipelago’s biodiversity faces ongoing threats from invasive species, including black and brown rats (*Rattus rattus, R. norvegicus*), house mice (*Mus musculus*), domestic cats (*Felis silvestris catus*), dogs (*Canis lupus familiaris*), cururu-toads (*Rhinella jimi*), and tegu lizards (*Salvator merianae*) (Micheletti et al. 2021).

### 2.2 Adaptive management protocol

Our RAM protocol (Figure 1) was based on applicable recommendations from Wegmann et al. (2011), Westgate et al. (2013) and Keitt et al. (2015), following the six key steps of AM (Westgate et al. 2013). The detailed protocol can be found in Appendix A. Modifications resulting from the RAM approach are detailed in the Results section.

**Figure 1.**
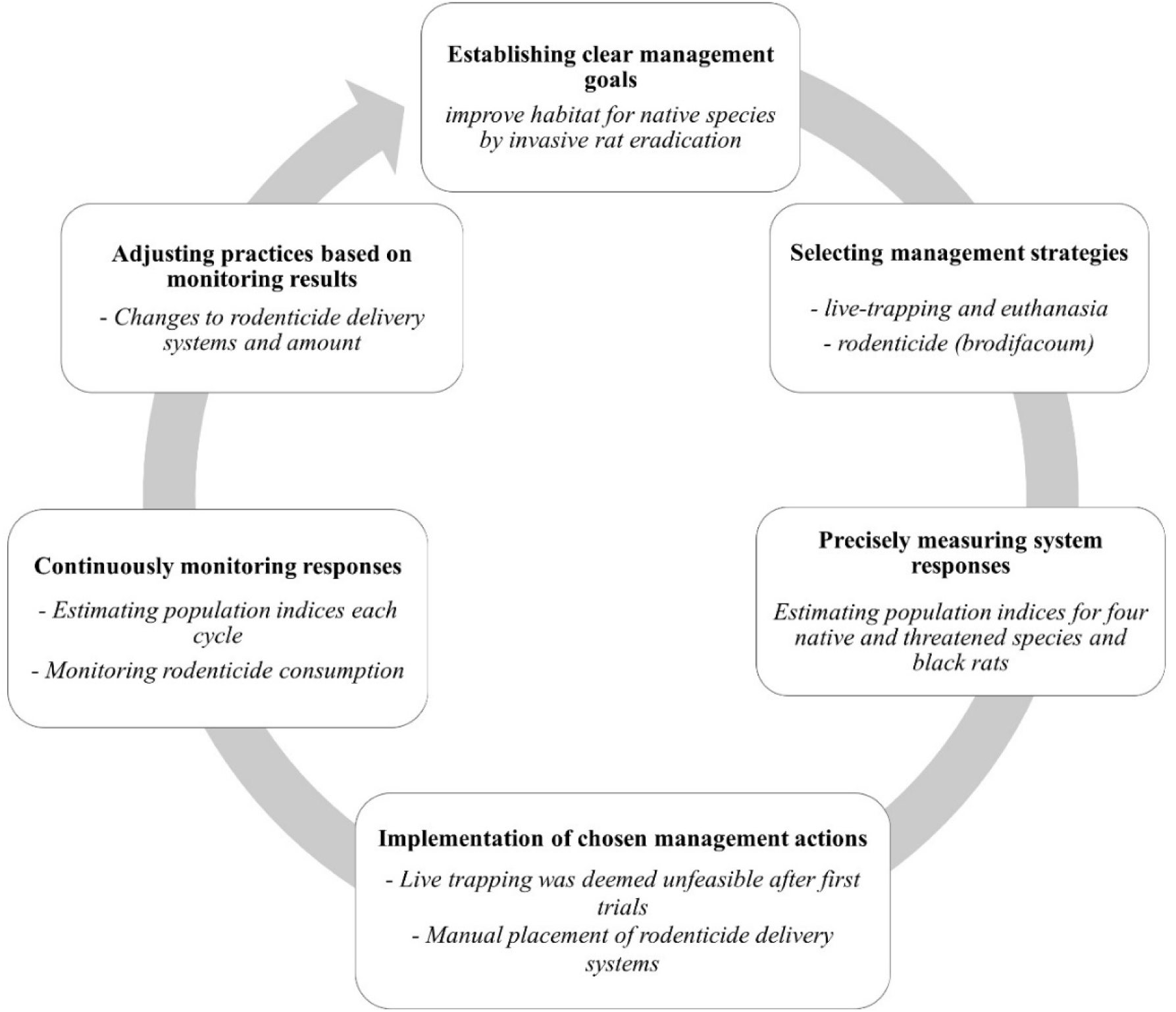
A scheme depicting our adaptive management protocol following the six iterative steps until objectives are met (modified from Westgate, 2013). In **bold the description** of the step, in *italic the specific actions* taken at each step.

#### 2.2.1 Establishment of Clear Management Goals

The primary goal was to improve habitat for the endangered insular land crab, endemic Noronha elaenia and mabuya, the threatened white-tailed tropicbird, and three booby species. While not monitored, the endemic Ridley’s worm lizard was also expected to benefit. To achieve this, we aimed to eradicate black rats, the primary threat, from Ilha do Meio. The project’s USD 60,000 budget was justified by its potential to prevent endemic species loss and protect seabird breeding grounds. This goal remained unchanged throughout the iterative adaptive management process.

#### 2.2.2 Selection of Management Strategies

We initially considered four management strategies: (1) live trapping followed by euthanasia, and rodenticide application via (2) aerial broadcast, (3) manual deployment, and (4) bait stations. All rodenticide strategies relied on brodifacoum (Table 1), chosen for its effectiveness and low failure rate (Parkes et al. 2011). Importantly, it is the only rodenticide authorized by governmental regulatory agencies for use in the archipelago. Preliminary calculations estimated that live trapping would require at least three months of daily effort with 20 traps (Russell et al. 2018), assuming similar rat densities to the main island, no trap-shy individuals, favorable weather, and no bait competition with land crabs. For rodenticide-based strategies, an estimated 9 kg of bait consumed by rats would be needed. After presenting the initial plan, discussions with funders and local managers revealed regulatory, logistical, financial, and ecological constraints, eliminating three of the four strategies. Therefore, only the manual placement of brodifacoum bait blocks in bait stations was approved for implementation.

**Table 1.**
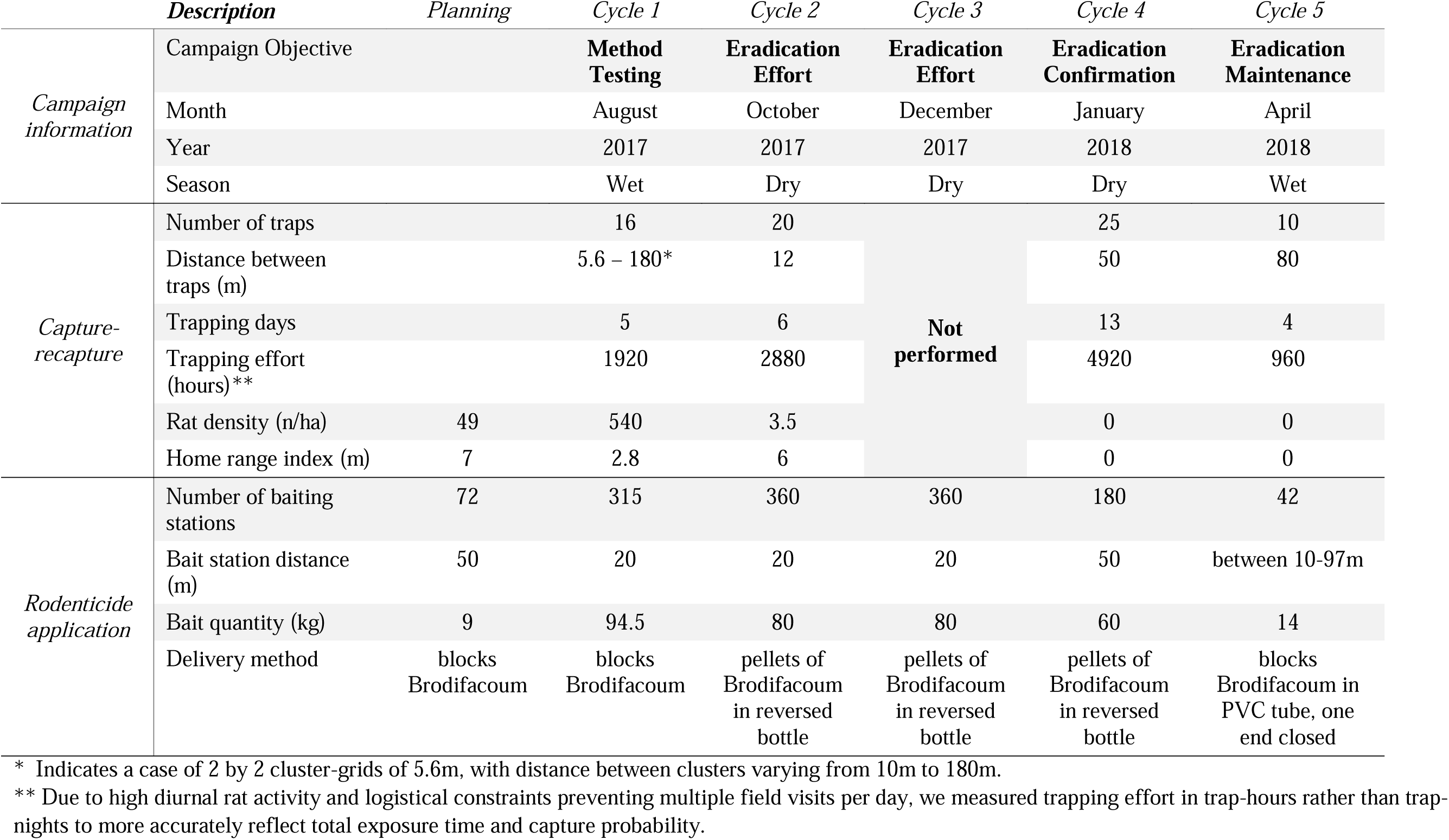
Summary of the iterative cycle information regarding the adaptive management for black rat eradication on the Ilha do Meio on the Fernando de Noronha archipelago (Brazil). Tomahawk galvanized wire trap, 450×210×210mm, foldable and with hooked-trigger were used for live-trapping. The bait used (KLERAT Mata Ratos, Syngenta Brasil) in block format weigh 20g, with approximately 1mg of rodenticide. Baited camera traps were deployed on 26^th^ May 2023 for 15 days, with a sampling effort of 1,080 hours, across all vegetation physiognomies, confirming eradication success.

#### 2.2.3 Measurement of System Responses

Habitat improvement was assessed using fixed-radius point counts for the Noronha elaenia, insular land crab, and mabuya, while seabird species were monitored through simple census surveys. Black rat density was estimated using spatially explicit capture-recapture (SECR) with 16 live traps (Table 2). Although 24 traps were planned, eight malfunctioned at deployment. Trap placement and numbers were adjusted based on preliminary results from Cycle 1. Detailed data and analysis methods are available in the manuscript’s repository (https://anonymous.4open.science/r/analysisIlhaDoMeio/README.md).

#### 2.2.4 Implementation of Management Actions

Baiting was initially planned for twice-weekly replenishment, with 72 stations placed 50 meters apart based on rat home-range estimates (Ringler et al. 2014; Robinson & Dick 2020). Rodenticide application was scheduled for the driest month (Samaniego-Herrera et al. 2014), with monitoring efforts aligned accordingly. All procedures adhered to animal welfare standards (CFMV 2012).

#### 2.2.5 Continuous Monitoring

The RAM framework was guided by two key purposes: assessing the benefits of black rat management on native species, and ensuring no negative impacts from rodenticide, following the precautionary principle. Initially, three iterations of rodenticide application and monitoring were planned to refine methods. Continuous monitoring included bait availability checks, wildlife monitoring, and black rat capture-recapture.

#### 2.2.6 Adjustment of Management Practices

Rodenticide application and black rat monitoring were adjusted in real time based on observations. Native species monitoring remained consistent but was extended for the Noronha elaenia due to new data from parallel projects. Eradication confirmation was initially planned two years after the last sighting using baited camera traps.

### 2.3 Native and Endemic Species Data Analysis

We used an open N-mixture population model (R package *unmarked*; Kellner et al. 2023) to analyze wildlife populations, testing 45 model formulations (Table A1). All model formulations are presented in Appendix B. Population growth predictions from the best models for Ilha do Meio were adjusted by subtracting those for Ilha Rata, creating the differential population growth index (DPGI) to isolate treatment effects from regional factors (e.g., extreme events, natural cycles, breeding seasonality). A linear model of DPGI over time was then fitted to assess native species’ population growth trends.

Originally, we planned to survey all three booby species and the tropicbird across both islands. However, the first expedition revealed that main nesting sites of the tropicbird, red-footed booby, and brown booby on Ilha Rata were inaccessible. Additionally, not all tropicbird nests on Ilha do Meio were reachable. Due to this lack of comparable control data, these species were excluded from modeling.

### 2.4 Hypothesis

We hypothesized that time since eradication started (TSE) would be a key predictor of native species’ population growth if rat eradication had a significant impact. We expected growth rates to increase over time, indicating positive effects on species recovery, and anticipated a corresponding positive trend in DPGI.

## 3 RESULTS

The RAM project had a total direct cost of approximately USD 60,000 (R$ 190,000 in 2017), or USD 3,300 per hectare. Each cycle followed three iterative steps: (1) rodenticide application and consumption monitoring, (2) continuous monitoring of black rat and native wildlife populations (3) management adjustments based on findings. Results are presented chronologically to illustrate the RAM process (Figure 1). A summary of the planning phase and five iterative cycles is presented in Table 1. Cycle-specific protocol adjustments, as well as wildlife and black rat monitoring results are detailed in Appendix C. Figure 2 summarizes native species DPGI and black rat density over time. All model objects and tables are available at https://zenodo.org/records/13830468.

**Figure 2.**
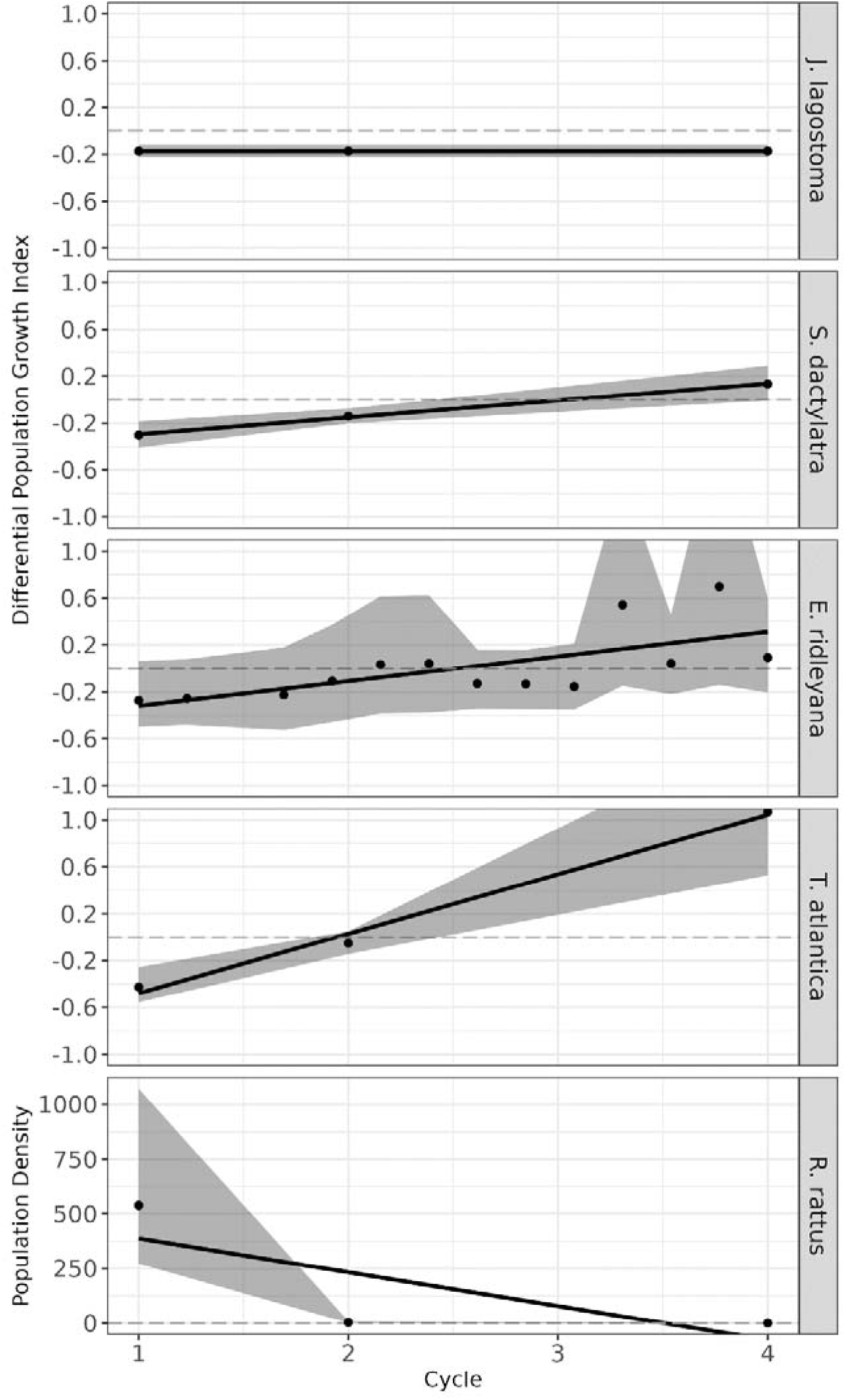
Differential population growth index across four cycles of real-time adaptive management for four native species (land crab, masked booby, Noronha elaenia and mabuya), as well as population density of black rat (*R. rattus*). The shaded areas represent the confidence intervals, with the solid lines depicting the population growth trends over time. Please note that the values for Noronha elaenia (*E. ridleyana*) were rescaled from 15 cycles to 4 due to data availability.

### 3.1 Management Cycles

In cycle 1 (August 2017), rodenticide baiting stations were deployed (Figure C1-A, Appendix C), and a capture-recapture trap grid revealed an unexpectedly high rat density (540 rats/ha). This prompted immediate adjustments: increasing rodenticide (+97 kg), adding bait stations (+315), and reducing station spacing (-30 m). Field observations indicated issues with mold and bait competition from crabs. In cycle 2 (October 2017), brodifacoum pellets were introduced in a reversed bottle system (Figure C1-B), and rat density dropped significantly (from 540 to 3.5 rats/ha). An additional 80 kg of brodifacoum was distributed across 360 bait stations. Cycle 3 (December 2017) focused solely on bait replenishment due to logistical constraints. By cycle 4 (January 2018), no rats were detected despite increased trapping efforts, leading to a 25% reduction in bait stations. This suggested eradication was achieved within five months. In cycle 5 (April 2018), efforts shifted to preventing reinvasion by reducing bait stations (n = 42) and establishing a *cordon sanitaire* with a new delivery method (Figure C1-C). No rats were photographed or captured after this, confirming eradication success.

### 3.2 Native and endemic species models and population parameters

The best models selected are presented in Appendix C (Table C1). Time since eradication started (TSE) significantly increased population growth for masked booby (0.14 ± 0.0380; p = 0.0002) and mabuya (0.46 ± 0.0971; p = 1.87e-06) on Ilha do Meio, with a positive but non-significant trend for Noronha elaenia (0.22 ± 0.1343; p = 0.0933). Positive population growth trends (DPGI) were observed for masked booby (0.14 ± 0.0059; p = 0.0263), mabuya (0.51 ± 0.047; p = 0.058), and Noronha elaenia (0.21 ± 0.069; p = 0.011), while land crabs showed no TSE effects or significant DPGI trends. These results highlight the role of eradication and environmental conditions in species recovery and stability. Population densities, detection probabilities, and growth rates for both islands based on the best models are presented in Table C2.

## 4 DISCUSSION

We demonstrate the successful application of real-time active adaptive management (RAM) to eradicate a highly dense black rat population (540 individuals/ha) on a tropical South Atlantic island. Eradication led to immediate biodiversity benefits, with rapid population growth observed in the endemic mabuya and Noronha elaenia, as well as the threatened masked booby. In contrast, land crab populations remained stable, unaffected by rat removal. To our knowledge, this is the first implementation of RAM for invasive species eradication in a tropical region, the first rat eradication on a Brazilian island, and the first detailed RAM case study applied to a biological system. By providing a comprehensive account of this process, we aim to offer a valuable reference for future conservation efforts.

### 4.1 Lessons learned about adaptive management and the RAM approach

AM has been widely discussed over the past half-century, but a persistent gap exists between theory and practice. Of 1,336 published AM studies, fewer than 1% included monitoring data (Westgate et al. 2013). Similarly, while 56% of evaluated studies (n = 187) advocated for AM, fewer than 5% reported successful implementation, with none quantifying benefits or costs (Rist et al. 2013). This trend extends to invasive species (IS) management, where fewer than 3% of 118 studies mentioning IS and AM provided implementation details (Appendix E), and only one incorporated real-time learning (Sciarretta et al. 2016), despite not achieving eradication. Importantly, none of these studies were conducted in the tropics, where eradication faces unique ecological and logistical challenges (Samaniego-Herrera et al. 2018; Samaniego et al. 2020).

A key barrier to AM adoption is the misunderstanding of its application Rist et al. (2013). The scientific definition of AM (Westgate et al. 2013) requires advanced scientific structuring and data analysis, skills often lacking among land managers—especially in underfunded tropical regions where conservation staff receive limited training (Passell 2000). This raises concerns about how scientific AM methods can be effectively implemented when managers lack the ability to assess success indicators. As an example, the Merri Creek revegetation study in Australia (McCarthy & Possingham 2007) exemplifies a textbook AM implementation, using Bayesian modeling and stochastic dynamic programming to optimize management strategies. Such approaches require specialized training and stable funding, conditions rarely met in tropical conservation programs. Until budget stability and manager training improve, AM will likely remain confined to well-funded regions.

Our RAM framework was effective in achieving eradication despite logistical, financial, ecological, and regulatory constraints, mainly due to a combination of the real-time iterative adjustments and tight transdisciplinary collaborations, as detailed below. The approach enabled complete eradication within five months while allowing for continuous monitoring of native species.

Regarding the direct results of the RAM approach, on Ilha Rata, land crab observations were positively influenced by rainfall (TR3) and negatively by moonlight (Hartnoll et al. 2006). Rats had no apparent impact on crab populations, which is supported by other studies (Gaiotto et al. 2020). While crabs consumed rodenticide, mortality risk remained low (Pain et al. 2000; Masuda et al. 2015), though secondary poisoning in humans warrants investigation, given possible ongoing poaching. Brodifacoum was undetected in crab claws and body tissues one month post-exposure but was present within the first month (Pain et al. 2000).

For masked boobies, rat eradication significantly boosted population growth on Ilha do Meio, as confirmed by DPGI results. On Ilha Rata, populations remained stable despite rats, consistent with studies showing booby persistence under predation (Bolton et al. 2011; Gouvêa & Mello 2017). However, evidence suggests rats disturb incubating adults, leading to nest abandonment (Priddel et al. 2005), a hypothesis supported by our findings. For mabuya, rat eradication strongly increased population growth (DPGI results). Models showed higher initial densities in treed areas, reflecting their habitat use. On the main island, rats consumed mabuya as 30.3% of their diet (Gaiotto et al. 2020), likely higher on Ilha do Meio due to the lack of human waste. Some mabuya ingested rodenticide, but monitoring showed no adverse effects. Further studies should confirm this. For the Noronha elaenia, population growth was negatively linked to TR3 due to its reliance on insect availability in the early rainy season (Licarião & De Brito Silva 2024) However, rat eradication significantly benefited population growth, as confirmed by DPGI trends. High variability in initial data suggested the need for supplementation, which was enabled by an independent monitoring project. While dispersal among islands is biologically possible, evidence suggests high site fidelity—it is found on only 3 of the 26 islands in the archipelago (Licarião & De Brito Silva 2024) Capture-recapture and genetic studies could confirm this.

Wildlife monitoring proved essential to monitor both the effects of the eradication and unwanted consequences of the management, detecting population trends by the fourth cycle. This underscores the value of real-time monitoring in conservation decisions.

### 4.2 Lessons learned about tight collaboration

The project’s success was heavily reliant on interdisciplinary collaboration between local environmental agencies, conservation practitioners, and scientific researchers. Close coordination facilitated rapid decision-making, particularly in adjusting strategies based on real-time data. This initiative was a pioneering collaboration between ICMBio, WWF-Brazil, and Instituto Tríade, marking a milestone in Marine Protected Area (MPA) management in Brazil. Historically, Brazil has lacked expertise in invasive species eradication (Sampaio & Schmidt 2013), with few large-scale management efforts (Silva & Alves 2011; Fonseca et al. 2024). Major invasive species threats remain unmanaged, such as the European wild boar (*Sus scrofa*)—one of the world’s 100 most invasive species (Lowe et al. 2000), now present across all six Brazilian biomes (Fonseca et al. 2024). Despite being targeted by the largest national invasive species control initiative, no effective management strategies have been implemented for wild boars (Kmetiuk et al. 2023). Our project highlights the power of interdisciplinary collaboration in AM projects (Dreiss et al. 2017), setting a precedent for future invasive species management in Brazil.

### 4.3 Lessons learned about handling the unexpected

Previous estimates for the main island of Fernando de Noronha suggested a rat density of 37 ± 12 rats/ha (Russell et al. 2018), but no data existed for Ilha do Meio. Surprisingly, our first-week estimate revealed an unprecedented 540 ± 195 rats/ha—11 times higher than the main island and exceeding all known records (64/ha in Hawaii; Tamarin & Malecha 1971; 66/ha in Mexico; Samaniego-Herrera et al. 2018; 187 rats/ha, 95% CI: 176-201; Vogt et al. 2014). This triggered real-time recalculations of rodenticide quantities, a challenge effectively addressed through the RAM approach. While this extreme density may raise questions, anecdotal evidence (e.g., high day and nighttime rat activity) long suggested an overpopulation, supported by high reproduction rates, abundant resources, and predator absence. Trap spacing of 25 to 50m has been suggested to ensure bait encounter for black rats (Ringler et al. 2014; Robinson & Dick 2020), though ideally, spacing should be less than twice the home range scale (σ) (Sun et al. 2014). Shorter distances (∼10m) improve density estimates in small areas (Sakamoto et al. 2015). Due to uncertainties, logistical constraints and trap malfunctions, we used an irregular grid, optimizing detection probabilities at the expense of precision—an approach recently supported for similar cases (Durbach et al. 2021; Freeman et al. 2022). While our model performed well, caution is needed when using irregular grids, as they may introduce biases (Smith et al. 2020).

Declaring eradication success posed another challenge, as no standardized protocol exists, and premature cessation has led to failures in past programs. Given uncertainties in detecting invasive species, eradication can rarely be guaranteed (Baker & Bode 2021), making ongoing surveillance essential (Ramsey et al. 2009, 2011; Samaniego Herrera et al. 2013). To that effect, we expanded trapping effort and conducted ad hoc monitoring of rodenticide consumption and indirect rat signs (e.g., bite marks, footprints, feces, predation on seabirds) during the fourth cycle. Although 15–20 trap-nights are often recommended for eradication confirmation (Russell et al. 2017), we set a conservative two-year window due to the significance of Brazil’s first rat eradication. The unexpected pandemic delayed confirmation until 2023, but despite concerns about reinvasion, Ilha do Meio remained rat-free five years post-eradication. While the island’s proximity to Ilha Rata is within black rat swimming ability (King et al. 1974; Shiels et al. 2014; Bagasara et al. 2016; Parkes et al. 2017), empirical evidence suggests no genetic flux between the islands (Gatto-Almeida et al. 2020).

### 4.4 Lessons learned about balancing social perception, environmental safety and efficiency

Although long-term rodent control is challenging, live-trapping followed by euthanasia has been effective for small areas (Duron et al. 2020) and is often preferred where rodenticides face social opposition (Duron et al. 2020; Fonseca et al. 2024). This option was considered but ultimately dismissed due to unlikely assumptions—including similar rat densities to the main island, absence of trap-shy individuals, stable weather, and minimal bait competition from land crabs.

While manual bait station placement was labor-intensive compared to aerial dispersal, strict regulations necessitated this approach. However, it had advantages: we used 18.25 kg/ha of rodenticide, aligning with temperate (Broome et al. 2014) and dry tropical islands (Samaniego-Herrera et al. 2018). Previous efforts on wet tropical islands have reported much higher bait usage (Wegmann et al. 2012; Samaniego-Herrera et al. 2018; Harper et al. 2019), demonstrating the variability and potential for substantially higher requirements in tropical regions (Pott et al. 2015). Yet, our work aligns with previous observations that rat eradication on some mesic-tropical islands may be achievable using typical temperate bait application rates (Griffiths et al. 2011).

Bait stations—while not the preferred approach among many practitioners—was designed within our precautionary RAM framework to detect potential negative impacts on native wildlife at the earliest stage. This was mandated by local authorities and especially critical given that no similar rodent management had ever been conducted on islands in Brazil, particularly within a national park. In the event of negative effects (e.g., declining population trends in monitored native species), our approach would allow for immediate intervention, including the potential removal of georeferenced rodenticide baiting stations if necessary. Additionally, reduced bait consumption by land crabs—likely due to improved bait delivery systems, which also mitigated mold issues during the wet season—ensured greater bait availability for rats, leading to effective lethal dosing across all rat territories (Howald et al. 2007).

### 4.5 Lessons learned about braiding policy and research

Our findings underscore the importance of designing adaptive strategies that account for both ecological and regulatory contexts (Walters & Holling 1990). The RAM framework proved effective in navigating these restrictions by integrating continuous monitoring and iterative adjustments (Williams 2011)—enabling real-time responses to emerging challenges. However, two key challenges must be addressed for long-term effective policy-making in conservation: (1) the lack of funding for research and awareness (Martin et al. 2012), and (2) the risk of excessive precaution reinforcing the status quo (Cooney 2005).

Without adequate financial support for parallel research, adaptive strategies may struggle to gain traction and are more likely to fail. In our project, not all researchers were employed full-time, and much of the work was conducted on a pro bono basis—an unfortunately common, yet unreported, situation in many developing countries. While funding may cover equipment, consumables, and occasionally technical staff, there is a chronic lack of support for compensating researchers involved in management projects. As a result, research often becomes a secondary objective in management initiatives, if it is considered at all. This issue requires urgent attention from funding bodies and society, as it may also contribute to the underrepresentation of AM projects in peer-reviewed literature, particularly in tropical regions (Martin et al. 2012).

To mitigate the reduced funding, we leveraged researchers’ field expertise in both data collection and rodent management, allowing these tasks to be carried out simultaneously. For instance, concurrent research and management enabled real-time coordination of bait replenishment with managers and field staff, ensuring bait availability until the next campaign based on observed consumption rates. This important integration of management and science (Williams 2011; Runge 2011) was crucial to the success of the RAM strategy, despite the reduced team size (six researches).

While stringent regulations are designed to mitigate risks to non-target species, they may also impose constraints that complicate the implementation of best practices in conservation, consequently allowing ecological degradation to persist (Cooney 2005). In our project, both safety and environmental concerns regarding the proximity of nesting seabird colonies (i.e., risk of accident) and the potential necessity of poison removal influenced the decision to forgo aerial baiting. Although aerial distribution has proven highly effective in numerous island eradication programs (e.g., Wegmann et al. 2012; Harper et al. 2019), a cautious, site-specific approach was deemed necessary to minimize unintended risks. Consequently, a labor-intensive manual baiting strategy was employed, increasing logistical complexity and potentially extending the duration of rodent pressure on wildlife before eradication was achieved (Russell & Broome 2016).

While precautionary measures aim to minimize risks to non-target species, they can also inadvertently hinder conservation efforts by delaying action and increasing the likelihood of eradication failure (Parkes et al. 2011). In our case, the decision to forgo aerial baiting in favor of a gradual, manual approach extended logistical complexity and potentially prolonged exposure risks before eradication was achieved. A more adaptive regulatory framework—one that integrates site-specific risk assessments and controlled trials—could help balance safety and environmental concerns with the urgency of invasive species management, ensuring effective and lower risk conservation outcomes.

The rapid recovery of habitat and biodiversity following rat eradication, as observed in our study, aligns with findings from other research on similar tropical islands (Miller-ter Kuile et al. 2021). This highlights the potential for significant ecological restoration in relatively short timeframes, reinforcing the importance of timely and decisive management actions.

### 4.6 Lessons learned about balancing risks

We acknowledge that our project carried risks. Following best practices (Broome et al. 2014; Keitt et al. 2015), would have required conducting trials and assessments before eradication, ensuring bait exposure to all rats within 1–3 days—a key factor in reducing failure risks. Manually placed bait stations, while necessary due to regulatory constraints, increased the risk of incomplete exposure, which could have led to failure. However, delaying eradication would have jeopardized native species, making immediate action the best option. To mitigate risks, we leveraged real-time monitoring and data science support, ultimately achieving successful eradication despite the odds.

Monitoring during eradication is generally discouraged due to potential confounding seasonal and environmental factors. However, it was essential for detecting adverse effects on native species and pausing interventions if necessary. By using Ilha Rata as a control site, we could isolate eradication effects, ensuring observed changes on Ilha do Meio were attributable to rat removal rather than natural fluctuations.

One major limitation was the lack of monitoring for rodenticide fallout, posing potential risks to marine mammals and other non-target fauna. While brodifacoum residues were undetected three years post-eradication at Palmyra Atoll (Wegmann et al. 2019), we strongly recommend future management actions secure funding to monitor potential environmental contamination and non-target impacts.

This study serves as a valuable example for conservation in highly regulated environments, demonstrating that invasive species eradication is achievable without compromising ecological or legal safeguards. However, our experience underscores the need for policy reform and greater investment in conservation to tackle underlying challenges. Collaboration between practitioners, policymakers, and researchers will be essential to ensuring the success of future eradication efforts, particularly in biodiversity-rich tropical regions.

## DECLARATION OF GENERATIVE AI AND AI-ASSISTED TECHNOLOGIES IN THE WRITING PROCESS

The authors used SCISPACE (https://typeset.io/) and ResearchRabbit (https://researchrabbitapp.com/home) for literature searches. SCISPACE queries included “Black Rat Eradication on Islands,” “Adaptive Management of Invasive Species,” and related terms. Manuscripts were selected based on title and abstract screening, similar to Google Scholar searches. To ensure comprehensive coverage, key references (Samaniego-Herrera et al., 2018; Westgate et al., 2013; Rist et al., 2013) were added to ResearchRabbit, allowing citation-based exploration. ChatGPT 4.0 was used for sentence refinement. All authors reviewed and edited the content and take full responsibility for its accuracy.

