## Appendix A for "Swiftly squeaky clean: lessons learned from eradicating an overpopulation of rats on an island of constraints"

Appendix A provides a detailed description of the real-time active adaptive management (RAM) protocol used in the black rat eradication project on Ilha do Meio, PE, Brazil (Figure A1). Our RAM protocol was based on applicable recommendations from Wegmann et al. (2011), Westgate et al. (2013) and Keitt et al. (2015), following the six key steps of AM (Westgate et al. 2013). It was also guided by two key purposes: assessing the benefits of black rat management on native species, and ensuring no negative impacts from rodenticide, following the precautionary principle. This appendix includes comprehensive information on the methods, from management goal establishment to precise implementation and adjustments of actions, offering a more granular perspective on the processes and decisions made. Modifications resulting from the RAM approach are summarized in the Results section of the main manuscript, and detailed in Appendix C. Overall, this section complements the manuscript by ensuring transparency and providing a valuable resource for replication in similar conservation efforts.

### 1. Establishment of clear management goals

The main management goal of our real-time active adaptive management (RAM) project was to improve the habitat for the endangered insular land crab (*Johngarthia lagostoma*), the endemics Noronha elaenia (*Elaenia ridleyana*) and mabuya (*Trachylepis atlantica*), , the threatened white-tailed tropicbird (*Phaethon lepturus*), and for the three booby species (*Sula dactylatra*, *Sula sula* and *Sula leucogaster*) that nest on Ilha do Meio (Figure A1). Although the endemic Ridley's worm lizard (*Amphisbaena ridley*) was not included in the project's monitoring efforts due to the complexity of its monitoring requirements, we anticipated that improved habitat conditions would benefit this species as well.

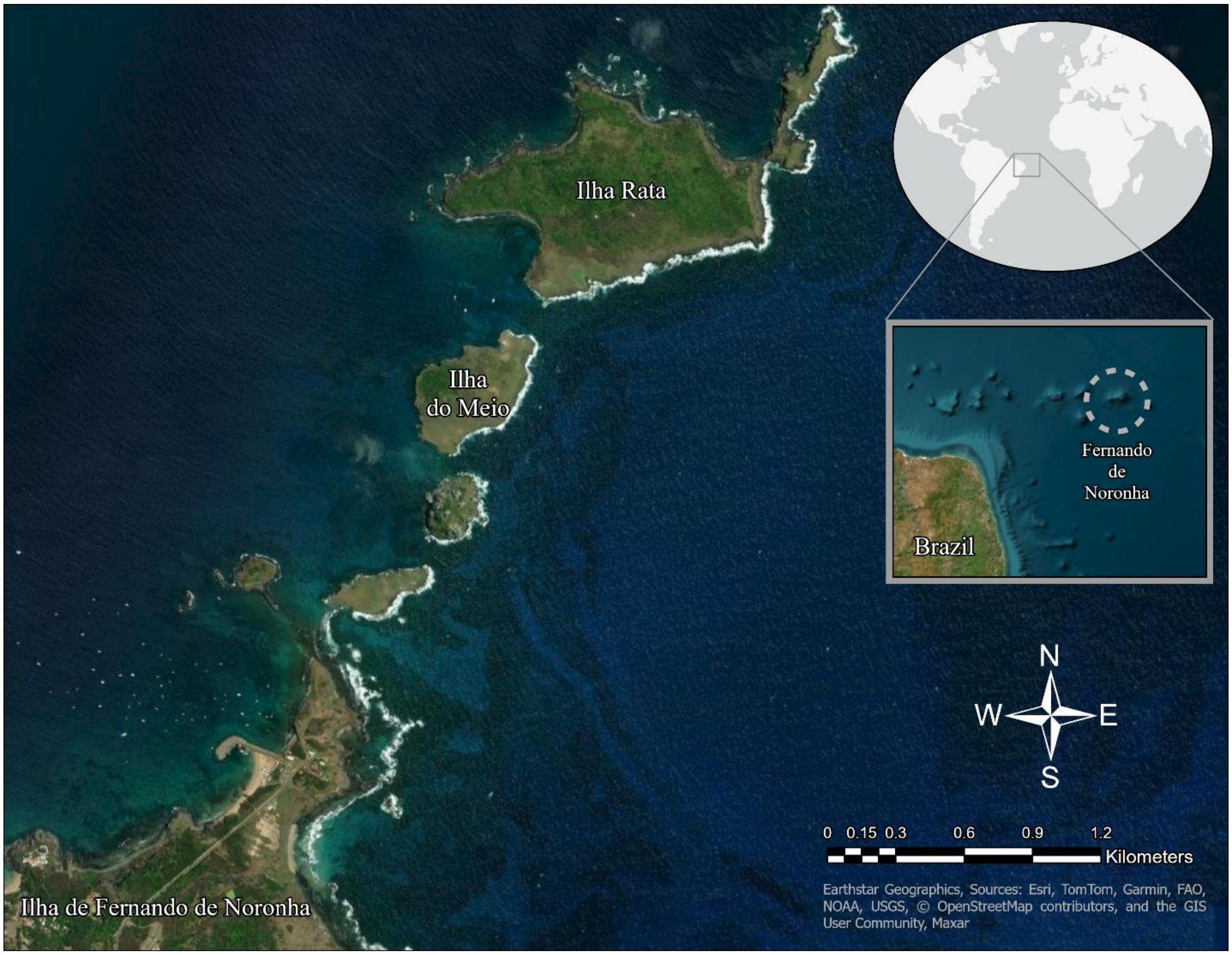


**Figure A1.** Study area composed of Ilha do Meio and Ilha Rata, pertaining to the archipelago of Fernando de Noronha and located north from the archipelago's main island with the same name. The archipelago is located off the northeastern coast of Brazil on the mid-Atlantic ridge.

To achieve our management goal, we aimed at eradicating black rats from the Ilha do Meio, the most importance threat to all native species present on the island. The benefits and costs of this RAM project to alternatives were empirically assessed during this planning phase. Rodent control – as opposed to eradication – was deemed not feasible due to the intricate logistics and long-term management commitment. Therefore, the only other alternative was identified as "no management" and its consequences were identified, in this order of importance, as the (i) potential loss of endemic species on the island, which are already present a restrict distribution, the (ii) reduction of breeding grounds for the sea bird species, and the (iii) maintenance of resource competition with the endangered insular land crab.

Although the third consequence was assessed as insufficient, the other two were weighted as worth it in relation to the potentially available project funding of USD 60,000. This goal was not changed during the RAM process.

### 2. *Selection of appropriate management strategies*

To eradicate the black rat population of the Ilha do Meio we selected two streams of management. The first stream focused on live-trapping of individuals, followed by euthanasia, which is often more socially accepted in the country than rodenticide usage (Fonseca et al. 2024). Previous studies on the main island of the archipelago have shown a capture rate of one rat per 48 hours at a 49 rats*$ha^{-1}$ density (Russell et al. 2018). This translates into an effort of at least three months with 20 traps. This strategy would require daily trips to the island to avoid mortality of trapped individuals by dehydration, and a larger team trained on euthanasia. Moreover, important assumptions would need to be made: rat density on Ilha do Meio as on the main island is very similar, absence of trap shy individuals on Ilha do Meio, no adverse weather influencing field logistics, no competition for baited traps between rats and land crabs.

The second stream focused on the use of rodenticides. We chose brodifacoum as our rodenticide, a vitamin K antagonist anticoagulant. Although brodifacoum is more toxic than other anticoagulant rodenticides, and as consequence, requires fewer feedings for a lethal dose, we chose it mainly due to its mechanism of action and the fact that it is the only rodenticide authorized for use in the archipelago. This rodenticide requires continuous ingestion and thus reduces negative effects on other species due to occasional ingestion. Brodifacoum has also presented significantly lower failure than other rodenticides on island eradications (Parkes et al. 2011), potentially due to its delayed effects; rats do not associate lethality with its ingestion, decreasing eradication failure rates. We had no means of monitoring the system for fallout. However, brodifacoum residues were not detected in the marine and terrestrial food web three years after rat eradication at Palmyra Atoll in Central Pacific (Wegmann et al. 2019), supporting our choice for this rodenticide. For this strategic stream, we primarily chose three possible rodenticide dispersion systems: aerial broadcast, manual dispersion of pellets, and manual dispersion of rodenticide in delivery systems (i.e., baiting stations).

Due to the lack of basic information on rodents for Ilha do Meio, we started from a black rat density of 49 individuals/ha. This value was the maximum observed density for the species on the main island of the archipelago (Russell et al. 2018). This prior density was needed for a first calculation of both the necessary number of live-traps and the amount of rodenticide to be used on the first application campaign.

To design the implementation schedule of the possible management actions the research team presented all four options, assumptions, and the preliminary cost calculations to funders and national park managers. During the plan assessment it was decided that the trapping followed by euthanasia presented too high of a risk due to the assumptions that needed to be made. Moreover, given the financial limitations and the environmental risks associated with helicopter deployment (i.e., potential need to remove rodenticide from the island should any monitored species present declining trends), manual placement of brodifacoum in delivery systems or bait stations (Appendix C, Figure 1C) was the only accepted method. This method has been demonstrated as an effective strategy for rat eradication on other islands (Keitt et al. 2015) and was well-suited to the size and field conditions of Ilha do Meio.

### *3. Precise measurement of system responses*

To monitor native wildlife, we used fixed-radius point count method (IPA; Hutto et al. 1986) for estimating population indices for the Noronha elaenia, the insular land crab and the mabuya. The counts were repeated over transects (i.e., existing trails) between 2 and 13 times per transect (n = 10 and 7, for Ilha do Meio and Ilha Rata, respectively), every 25m for mabuya and land crab and every 60m for Noronha elaenia. Counts were performed on both the Ilha do Meio and Ilha Rata, the last one being used to control for seasonal variation on indices. Transects were also performed at approximately the same time of the day, between 6 am and 8 am for crabs and avifauna, and between 9 am and 11 am for the mabuya. We also used a simple census survey for the number of *Sula* individuals in breeding sites across the islands. Although counts may be biased by land cover type and bird movement (Buckland et al. 2008), the bird's nesting site during the study presented relative low vegetation coverage, allowing for a 'snapshot' of the seabird population. During the breeding season, adults actively defend their nests, which limited bird movement, facilitating counts.

To measure the response of the RAM for black rat eradication on Ilha do Meio, we estimated black rat density using spatially-explicit capture–recapture (SECR) with the R package `secr` (Efford 2023) during monitoring campaigns. These monitoring campaigns happened concomitantly with rodenticide application campaigns, and were planned at first to last 5 and 10 days, respectively. SECR was used with a half-normal detection curve. We used at first 16 traps (Tomahawk galvanized wire trap, 450x210x210mm, foldable, trigger with hook) placed as 2 by 2 grids of 5.6m, with distance between clusters varying from 10m to 180m. Although we had initially planned using 24 traps, eight of them malfunctioned at deployment. Moreover, each trapping sampling session data (i.e., campaign data of consecutive days) was used as a stand-alone, independent dataset. This was done due to the adjustments needed on number and disposition of traps for each new monitoring campaign, as described below. Preliminary model tests (i.e., model comparison using AIC) supported the assumption of lack of both sex and landscape effects on capture probability and density, even though in some island patches the vegetation varies from shrubby to arboreal. We used the island's contour as the natural analysis boundary, removing the need to define a buffer for the traps, which can impact the analysis (Efford 2023). All data and detailed analysis code can be found in the manuscript's repository (https://anonymous.4open.science/r/analysisIlhaDoMeio/README.md).

### 4. *Implementation of chosen management actions*

A conservative estimate for acute oral $LD_{50}$ (i.e., the lethal dose for 50% of the individuals) of brodifacoum for *R. rattus* has been shown to be 0.77 mg/kg of body mass (Mathur & Prakash 1981). Using a linear relationship between dose and lethality, we calculated that a lethal dose ($LD_{100}$) of brodifacoum for any individuals to be 1.54mg/kg of body mass. While a linear relationship is unlikely to be an accurate method for calculating $LD_{100}$, it is the best approximation in our case due to lack of information on $LD_{100}$ for the black rat population on the archipelago. Black rats have been shown to weight a maximum of 305g under controlled environment with food being provided *ad libitum* (Bentley & Taylor 1965). The lethal dose of brodifacoum per individual was then preliminarily calculated as 0.5mg. Our brodifacoum first bait of choice (KLERAT, Syngenta) contained approximately 20g per wax block, with the rodenticide accounting for approximately 1mg. This means that each rat eating approximately half (10g) of a wax block would ingest a lethal dose of the rodenticide. Although calculating the individual ingestion rate is not the standard management practice – normally managers simply ensure sure that all rats are exposed to poison for minimum period of time – our approach focused on both avoiding excessive acquisition of rodenticide, as well as excessive dispersal of rodenticide on the island, reducing ecological risks.

Based on the best information available for the rat density (maximum of 49 individuals/ha; Russell et al. 2018) and the total island area of 18ha, we calculated the total population on the island to be 882 individuals. With a block needed for each two individuals, we calculated 441 blocks (approximately 8.82kg) would suffice to eradicate the black rat from the Ilha do Meio, provided all bait was consumed by all rats, exclusively, within ten days (Bhat & Sujatha 1989). Based on logistics, we opted for the bait administration to be done twice within a week, with 3 to 4 blocks per station, lasting 3 to 4 days with the consumption being calculated as about 3g of bait per individual per day. This resulted in 72 baiting stations with a distance of approximately 50m between each station. Blocks were protected by a plastic cover to avoid rain and possible seabird interaction with the rodenticide (Appendix

C, Figure 1C, Panel A). The first rodenticide application campaign was planned to be carried out in tandem with rodent trapping and wildlife monitoring, leveraging the daily presence of researchers on the island for five days in a row to check the bait stations. Rodenticide, rodent trapping and wildlife monitoring campaigns were planned for about every two months. Capture and handling techniques, as well as rodenticide usage conformed to high standards of animal welfare (CFMV 2012).

### 5. Continuous monitoring of system responses

Originally, three iterations of rodenticide application and population monitoring were planned: the first allowing for adjustments in the methods based on the collected new information, the second focusing on the eradication of the black rat population, and the third for confirmation of eradication success. The continuous monitoring of system responses consisted of three tasks: (1) monitoring stations for bait availability, (2) performing population counts for native wildlife, and (3) performing capture-recapture of black rats for population monitoring.

### 6. Adjustment of management practice in response to monitoring results

As the crucial point of RAM, we made method adjustments to black rat's rodenticide application, not only between campaigns, but also within them. Wildlife population monitoring method remained the same through the entire study. However, for the Noronha elaenia, monitoring efforts were maintained mostly twice a year (i.e., during the COVID pandemic, May 2019 until September 2021 this was not performed), until 2023 as data was available from a parallel monitoring project. Regarding black rat's population monitoring, the number of traps, distance between them, trapping effort and trap's spatial distribution on the island were adapted between campaigns following a careful review of population estimate results. During black rat monitoring activities, researchers also checked baiting stations regularly. This allowed for a real-time coordination for bait replenishment with managers and field staff as to ensure bait availability both within and between campaigns, based on observed daily bait consumption. Rodenticide quantity and as well as number of bait stations were recalculated within campaigns, at the end of each trapping session. The rodenticide delivery method was adapted between campaigns using field observations, bait consumption and black rat population information. We planned on confirming rat eradication after two years of the last sighting, by baiting camera traps.

### 7. *Native and endemic species' data analysis*

Originally, we planned to survey all three booby species and the tropicbird across both islands. However, the first expedition revealed that nesting sites of the tropicbird, red-footed booby, and brown booby on Ilha Rata were inaccessible. Additionally, not all tropicbird nests on Ilha do Meio were reachable. Due to this lack of comparable control data, these species were excluded from modeling. To model all wildlife species for which we managed to collect data on both islands, we used an open N-mixture population model (Dail & Madsen 2011), which is available in the R package `unmarked` (Kellner et al. 2023). Different model formulations (n = 45) were tested for all species (Table A1) and are shown in Appendix B. For Noronha elaenia a robust design model was fitted, while immigration was not considered in the model for any of the species due to physical isolation (i.e., mabuya) or site fidelity (i.e., land crab, Noronha elaenia, masked booby).

**Table A1.** Model formulations with tested variables on each model parameter for all four endangered – land crab (*Johngarthia lagostoma*) and masked booby (*Sula dactylatra*) – and endemic - Noronha elaenia (*Elaenia ridleyana*) and mabuya (*Trachylepis atlantica*) – species monitored on Ilha do Meio and Ilha Rata (Fernando de Noronha, Brazil) during a black rat (*Rattus rattus*) eradication performed in 2017/2018 using rodenticide in baiting stations (brodifacoum).

| **Species** | **Parameter name** | **Parameter representation** | **Land Cover** | **Observer ID** | **Moon Phase** | **TR3*** | **TSE**** |
|---|---|---|---|---|---|---|---|
| *Johngarthia lagostoma (land crab)* | lambda | Initial population | ✓ | | | | |
| | gamma | Population growth | | | | | ✓ |
| | det | Detection | | | ✓ | ✓ | |
| *Sula dactylatra (masked booby)* | lambda | Initial population | | | | | |
| | gamma | Population growth | | | | | ✓ |
| | det | Detection | | | | | |
| *Elaenia ridleyana (Noronha elaenia)* | lambda | Initial population | | | | | |
| | gamma | Population growth | | | | ✓ | ✓ |
| | det | Detection | | | | | |
| *Trachylepis atlantica (mabuya)* | lambda | Initial population | ✓ | | | | |
| | gamma | Population growth | | | | | ✓ |
| | det | Detection | | ✓ | | | |

* Total rainfall in the previous three months
** Time since eradication started

The best model formulation for each species and island (i.e., Ilha Rata as control, Ilha do Meio as treatment) was chosen based on Akaike's Information Criterion (AIC; Akaike 1992). The model fitted using the function `pcountOpen` allows for poisson (P), negative binomial (NB) or zero-inflated (ZIP) distributions. For each species on each island (i.e., treatment and control), the null formulation of each one of these three models was tested, identifying which model presented significant zero-inflation or dispersion, or none of these, suggesting the use of ZIP, NB and P, respectively. Zero inflation was also tested by checking the proportion of real over expected zeros using the JVDB score test (Van Den Broek 1995) and the number of zeros over the entire dataset not surpassing 20%. A ZIP distribution was chosen if significancy was reached either on the zero-inflation parameter of the null ZIP model, or on the JVDB score test and percentage. In case both zero-inflation and overdispersion were present, the distribution was chosen based on the best AIC between the ZIP and the NB null models.

To select an appropriate carrying capacity (K) for the model, representing the upper limit of individuals that can be modeled or the maximum expected population size, we iteratively tested the best model formulation with increasing K values. This process continued until the Akaike Information Criterion (AIC) stabilized, with a tolerance of less than 1, ensuring relative stable parameter estimates. Tolerance was implemented as an important tradeoff exists between with increases in K and computation time and power. Best model estimates were then used to predict population growth through time (i.e., ecologically known as lambda, but referred to as gamma in unmarked). Then, these were adjusted by subtracting equivalent predictions from Ilha Rata. This subtraction yielded a “Differential Population Growth Index” (DPGI), which serves to isolate the specific treatment effects from broader regional influences (e.g., extreme weather events, natural population cycles, or breeding seasonality). Using this refined metric, a linear model of DPGI over time was fitted to evaluate how native species’ population growth trajectories responded to the management actions on Ilha do Meio.
