## Appendix C for "Swiftly squeaky clean: lessons learned from eradicating an overpopulation of rats on an island of constraints"

Appendix C provides a detailed account of the changes made to the real-time active adaptive management (RAM) protocol over five cycles during the black rat eradication project on Ilha do Meio, PE, Brazil. The appendix outlines the adjustments made to rodenticide application, bait station setup, and trapping efforts in response to real-time data and field observations. Each cycle details key findings, such as recalculated rat densities, changes in baiting strategies, and the evolving effectiveness of the rodenticide delivery systems. The RAM approach incurred a total direct cost of around USD 60,000 (R$ 190,000 in 2017), equivalent to approximately USD 3,300 per hectare. Each iteration of the management cycle proceeded through three key steps: (1) applying rodenticide and monitoring its consumption, (2) continuously monitoring black rat and native wildlife populations, (3) adjusting management strategies based on the collected data. All model objects and tables are available at https://zenodo.org/records/13830468.

### 1.1. Cycle 1

During the first real-time monitoring cycle (August 2017), we started the setup of the rodenticide baiting stations (Figure C1, Panel A) while setting up the capture-recapture trap grid. At the end of the first week, rat density was estimated to be on average 11 times higher with a smaller home range than the estimated on the main island (540 ± 195 ind/ha and 2.8 m home range). This prompted an immediate recalculation of rodenticide needed and the number of stations to be used. The updated amount of rodenticide was then calculated as approximately 97 kg (0.01 kg per individual * 540 individuals/ha * 18 ha), and the shortfall was communicated to managers for immediate purchase of brodifacoum.

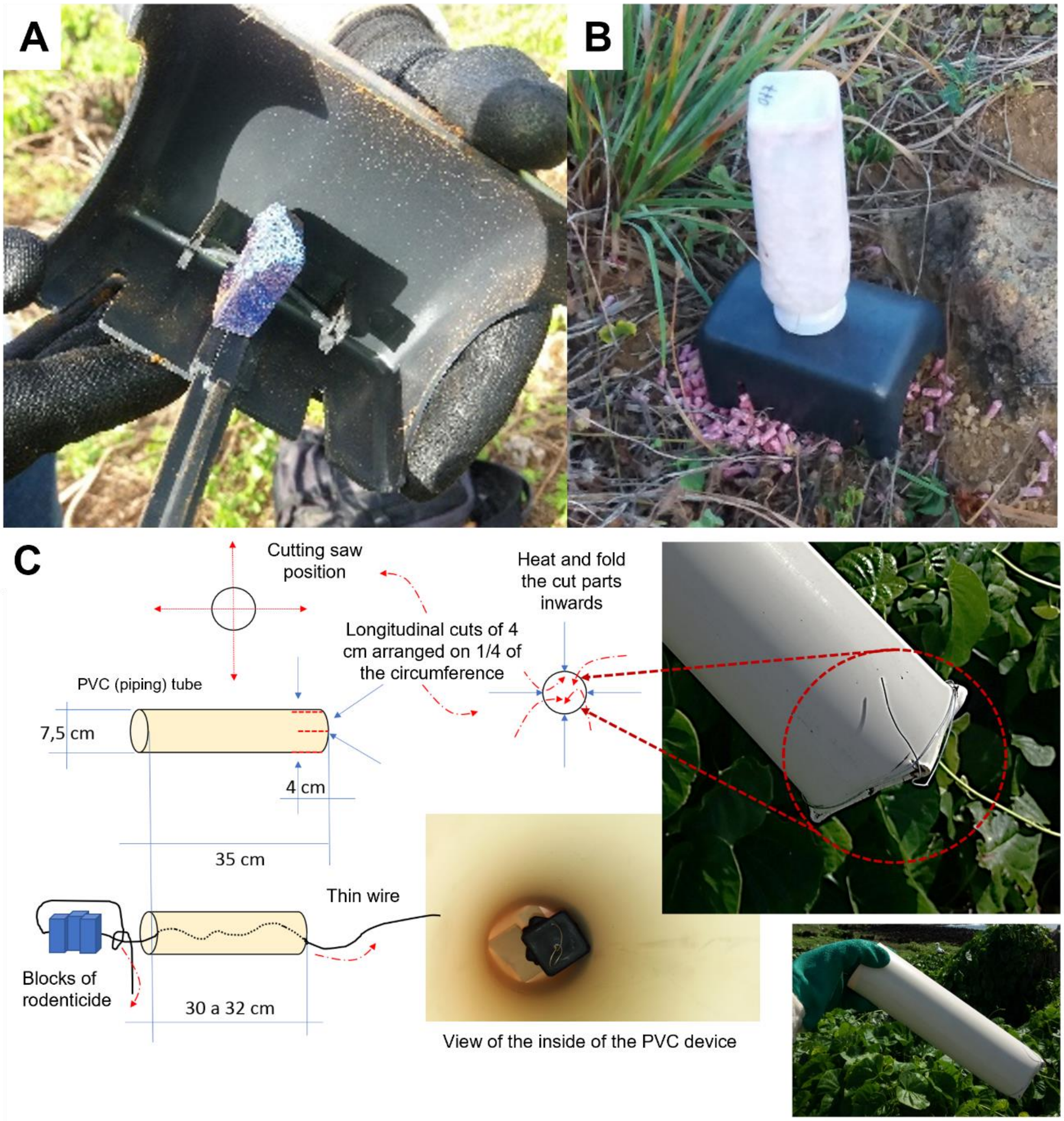


**Figure C1.** Blocks of rodenticide (brodifacoum) were protected by a plastic cover to avoid rain and seabird consumption (A) and pellets and stored theses in a reversed bottle system (B). A mechanism was also built to compose the sanitaire cordon, as a means of helping prevent reinfestation (C) and also avoiding bait consumptions by land crabs.

The number of baiting stations as well as the distance between these were also recalculated, increasing the bait ratio per station by approximately 60%. At the end of the first cycle, there were 315 baiting stations spread approximately 20m apart over the entire island. During the campaign, researchers observed bait blocks being taken in full by rats and crabs. Moreover, when checking for bait availability between the first two cycles, it was observed that some of the bait blocks had become moldy, likely reducing its palatability. These observations could indicate a that homogenous intake of the rodenticide by the whole population could be jeopardized, which prompted a discussion about changes in the delivery method. Moreover, during the first expedition the nesting sites for red-footed and brown booby on Ilha Rata were found to be inaccessible. Therefore, although our data has counts for these species for Ilha do Meio, we did not model them due to the lack of control (i.e., Ilha Rata) information. Due to inaccessible nests on both islands, data on tropicbirds was not collected.

#### 1.2. Cycle 2

Field observations from the first cycle resulted in a decision to change the rodenticide delivery method for the second cycle (October 2017). We exchanged brodifacoum blocks for pellets and stored 220 g of theses in a reversed bottle system (Figure C1, Panel B) to ensure a more homogeneous and longer rodenticide delivery to the black rat population. Expecting a slow decrease in the rodent population due to food availability and competition for rodenticide by land crabs, another 80 kg of brodifacoum were acquired between the two cycles. The results of the capture-recapture analysis from the first cycle also suggested that our effective sampling area (*esa*) was smaller than expected (574 $m^2$; Table C1), potentially due to our irregular grid. However, to our knowledge, no studies have explored irregular grid effects coupled with open N-mixture models of SECR, leaving questions about how to accurately estimate effective sampling areas in these scenarios. Yet, to decrease uncertainty in black rats' estimation, we increased the trapping effort to 2880 trapping hours (six trapping days), as well as the number, and changed the distance between the traps to 20 units and a uniform 12 m, respectively. Capture-recapture during the second cycle showed a 32-fold decrease in black rat abundance, from 540 to 3.5 individuals per ha. Although, about 3 kg of brodifacoum could suffice for the newly calculated population size, we opted to still use the purchased brodifacoum with the adapted delivery method. This ensured a more homogenous delivery across the island, over 360 baiting stations. The 45 new baiting stations were set more densely on the arboreal patch on the island, where the higher food availability observed might have supported higher rodent density.

#### 1.3. Cycle 3

Due to extreme weather, an important logistic constraint in our study site, no rodent or wildlife monitoring were performed as planned in the third cycle (December 2017). Nonetheless, to avoid losing the effort on rodent suppression, rodenticide stations were replaced by fresh rodenticide, maintaining the 360 baiting stations and repeating the amount of 80 kg distributed across these. During the third cycle a reduction on the number of stations was planned to take place in the fourth cycle, based on the lack of rodent activity (i.e., indirect rodent signs like chewing marks, feces and direct observations) in opportunistic field observations.

#### 1.4. Cycle 4

During the fourth cycle (January 2018) the approximately 220 g of bait per delivery station was kept. This resulted in a bait usage of approximately 60 kg. Rodent trapping effort was increased to 25 traps, placed approximately 50 m apart, but covering all land cover physiognomies (i.e., arboreal, shrubby and open). Trapping activities lasted 13 days during this cycle, with the single objective of detecting presence of rodents, not anymore estimating population size. Even with an effort of 4,920 trapping hours, no rodents were captured, suggesting the black rat population had been potentially successfully eradicated. Therefore, the number of traps was reduced by 25%, keeping an average distance between baiting stations of 50 m.

### 1.5. Cycle 5

Due to the potential black rat eradication confirmed in the fourth cycle, it was decided that the fifth (April 2018) would be the last cycle of the real-time iterative management. This cycle's objective differed from previous ones by focusing on establishing a protective ‘cordon sanitaire’ (i.e., trap barrier) along the island's perimeter and at key strategic points within it to prevent rat reinvasion. Therefore, the number of baiting stations was reduced to 42 with a median of 38 m, and a maximum of 83 m between these. About 9.3 kg of rodenticide was distributed across the baiting stations. Bait delivery system was again adapted to the new objective. Brodifacoum blocks were tied with a metal string to a closed side of a 3-inch PVC tube, allowing for black rats to come in and feed on the block, but securing it to the tube to avoid land crab consumption (Figure C1, Panel C).

Capture-recapture was also performed in this cycle, but with reduced effort. Across the island, 10 traps were set for 4 nights, approximately 80 m apart. This totaled 960 trapping hours and resulted in no captures. At the end of cycle 5, it was decided that the project had been successfully concluded. Eradication confirmation was planned to be announced in five years’ time after camera-trap effort had not registered further rat activities. Three camera-traps (Bushnell Thropy Cam HD – Essential E3 Brown, model 119837) were installed on the island on 26th May 2023 for 15 days, totaling 1080 hours of sampling effort across all vegetation physiognomies. There were no records of rats during this expedition.

### 1.6. Native and endemic species models and population parameters

The best models selected are presented in Table C2. Regarding the threatened species, for land crabs the best model for Ilha Rata presented a non-significant effect of total rainfall in the previous three months (TR3; 0.13 ± 0.073; p = 0.0742) and a significant negative effect of moon phase (i.e., towards new moon; -0.173 ± 0.068; p = 0.0109) on observation (*p*). Landscape cover affected initial population (*lambda*) for Ilha do Meio (i.e., positive in treed areas; 1.32 ± 0.1826; p = 5.55e-13), and we also observed a significant positive effect of TR3 (0.73 ± 0.060; p = 9.23e-34) and a significant negative effect of moon phase (i.e., towards new moon; -0.39 ± 0.070; p = 1.60e-08). For this species, TSE was not in the best model. For the masked booby, the only model chosen for Ilha Rata, our control, was a constant model. For Ilha do Meio, our treatment island, the best model presented TSE as a covariate for population growth (*gamma*). Population growth estimate for Ilha do Meio was significantly positively influenced by TSE (0.14 ± 0.0380; p = 0.0002).

Regarding the endemic species, the model for Noronha elaenia for Ilha Rata, presented a significant negative effect of TR3 on population growth (-0.33 ± 0.1485; p = 0.0253), while for Ilha do Meio, TR3 and TSE had a significant negative (-0.49 ± 0.1563; p = 0.0017) and a positive (0.22 ± 0.1343, p = 0.0933) effect on population growth, respectively. Lastly, the best model selected for mabuya for Ilha Rata showed an influence of observer on observation for both islands. For Ilha do Meio, landscape cover affected initial population (i.e., positive in treed areas; 1.09 ± 0.1536; p = 1.16e-12) and TSE had a significant positive effect on population growth (0.46 ± 0.0971; p = 1.87e-06). Covariate estimates, standard errors, z and p values, as well as AIC, model mixture and K for the best models are presented in Table 5.

While land crab presented a constant fitted linear model of DPGI through time (1.37e-17 ± 1.19e-17; p = 0.454), masked booby (0.14 ± 0.0059; p = 0.0263), mabuya (0.51 ± 0.047; p = 0.058) and Noronha elaenia (0.21 ± 0.069; p = 0.011) presented positive trends. Native species' population densities, observation probabilities (i.e., detection) and population growth rate for both islands for the best models are presented in Table C3.

**Table C1.** Spatially explicit capture-recapture parameters of black rats (*Rattus rattus*) on the Ilha do Meio (Brazil) during the first (cycle 1) and the second (cycle 2) rodenticide application campaigns, which happened in August and October 2017. Parameters estimated are effective sampling area (esa), density (D) in hectares, detectability (g0), index of home range size (sigma). For density estimates, standard error of estimate (SE), lower (LCL) and upper (UCL) confidence limits, coefficient of variation for population density (CVn), coefficient of variation of individual detection (Cva), and coefficient of variation in detection (CVD) were also calculated. For the detection parameters (g0 and sigma), the respective link function is described. During cycle 3 only rodenticide administration took place, and during cycles 4 and 5 there were no black rat captures.

| Cycle | Parameter | Estimate | SE | LCL | UCL | CVn | Cva | CVD | Link |
|---|---|---|---|---|---|---|---|---|---|
| 1 | esa | 0.05740268 | | | | | | | |
| | D | 540.0444614 | 194.9238 | 271.9902 | 1072.274 | 0.1796 | 0.3130 | 0.3609 | |
| | g0 | 0.1186521 | 0.04032339 | 0.059468 | 0.222787 | | | | logit |
| | sigma | 3.6021484 | 0.59044688 | 2.617945 | 4.956358 | | | | log |
| 2 | esa | 6.041693 | | | | | | | |
| | D | 3.475847 | 1.706929 | 1.397699 | 8.64386 | 0.2182 | 0.4399 | 0.4910 | |
| | g0 | 0.09114396 | 0.02956253 | 0.04746145 | 0.1679425 | | | | logit |
| | sigma | 58.96512987 | 21.13140513 | 29.83429736 | 116.5399171 | | | | log |

**Table C2.** Covariate estimates, standard errors, z and p values, as well as AIC, model mixture (poisson, negative binomial or zero inflation) and K (maximum number of individuals) for the best models chosen for each one of the endangered or threatened species during an eradication attempt on Ilha do Meio (Fernando de Noronha, Brazil).

| species | island | model | parameter | covariate | estimate | Standard Error | z | p | AIC | Model Mixture | K |
|---|---|---|---|---|---|---|---|---|---|---|---|
| *Trachylepis atlantica* | Ilha do Meio | lam(landscapeCover)<br>gamma(TSE)<br>p(observerID)<br>iota(.) | lambda | (Intercept) | 4.634611 | 0.365236 | 12.68937 | 6.77E-37 | 587.7751 | ZIP | 650 |
| | | | lambda | landscapeCoveropen | -3.53705 | 0.8364 | -4.22889 | 2.35E-05 | 587.7751 | ZIP | 650 |
| | | | lambda | landscapeCoversemi-open | -2.14083 | 0.818179 | -2.61658 | 0.008882 | 587.7751 | ZIP | 650 |
| | | | lambda | landscapeCovertreed | 1.092222 | 0.153612 | 7.110251 | 1.16E-12 | 587.7751 | ZIP | 650 |
| | | | gamma | (Intercept) | 0.00922 | 0.049946 | 0.184599 | 0.853544 | 587.7751 | ZIP | 650 |
| | | | gamma | scale(TSE) | 0.462656 | 0.09706 | 4.766691 | 1.87E-06 | 587.7751 | ZIP | 650 |
| | | | det | (Intercept) | -2.47652 | 0.795474 | -3.11326 | 0.00185 | 587.7751 | ZIP | 650 |
| | | | det | observerIDMangini | -1.05018 | 0.881041 | -1.19198 | 0.233269 | 587.7751 | ZIP | 650 |
| | | | det | observerIDVerona | -2.14182 | 0.883423 | -2.42445 | 0.015331 | 587.7751 | ZIP | 650 |
| | | | det | observerIDVini | -1.4007 | 0.847839 | -1.65208 | 0.098519 | 587.7751 | ZIP | 650 |
| | | | psi | psi | -12.9792 | 199.2237 | -0.06515 | 0.948055 | 587.7751 | ZIP | 650 |
| | Ilha Rata | lam(.)<br>gamma(.)<br>p(observerID)<br>iota(.) | lambda | (Intercept) | 5.19913 | 2.708121 | 1.919829 | 0.05488 | 297.7525 | ZIP | 450 |
| | | | gamma | (Intercept) | -0.00704 | 0.080663 | -0.08725 | 0.930474 | 297.7525 | ZIP | 450 |
| | | | det | (Intercept) | -5.23503 | 2.730988 | -1.9169 | 0.055251 | 297.7525 | ZIP | 450 |
| | | | det | observerIDMangini | 0.739664 | 0.309288 | 2.391508 | 0.016779 | 297.7525 | ZIP | 450 |
| | | | det | observerIDVini | 0.768243 | 0.407786 | 1.883938 | 0.059573 | 297.7525 | ZIP | 450 |
| | | | psi | psi | -5.62101 | 3.729429 | -1.50721 | 0.131758 | 297.7525 | ZIP | 450 |
| *Elaenia ridleyana* | Ilha do Meio | lam(.)<br>gamma(totalRainfallThreeMonths+TSE)<br>p(.)<br>iota(.) | lambda | (Intercept) | 1.599695 | 0.488569 | 3.274243 | 0.001059 | 424.1632 | ZIP | 250 |
| | | | gamma | (Intercept) | -0.01973 | 0.118374 | -0.16666 | 0.867636 | 424.1632 | ZIP | 250 |
| | | | gamma | scale(totalRainfallThreeMonths) | -0.48951 | 0.156321 | -3.13146 | 0.001739 | 424.1632 | ZIP | 250 |
| | | | gamma | scale(TSE) | 0.225422 | 0.1343 | 1.678497 | 0.09325 | 424.1632 | ZIP | 250 |
| | | | det | (Intercept) | -0.03968 | 0.203835 | -0.19467 | 0.845653 | 424.1632 | ZIP | 250 |
| | | | psi | psi | -5.3502 | 14.58272 | -0.36689 | 0.713704 | 424.1632 | ZIP | 250 |
| | Ilha Rata | lam(.)<br>gamma(totalRainfallThreeMonths)<br>p(.) | lambda | (Intercept) | 1.129187 | 0.577491 | 1.955331 | 0.050544 | 423.8833 | P | 250 |
| | | | gamma | (Intercept) | 0.017988 | 0.128122 | 0.140399 | 0.888345 | 423.8833 | P | 250 |

| | | | | | | | | | | | |
|---|---|---|---|---|---|---|---|---|---|---|---|
| | | iota(.) | gamma | scale(totalRainfallThreeMonths) | -0.33213 | 0.148481 | -2.23683 | 0.025298 | 423.8833 | P | 250 |
| | | | det | (Intercept) | 0.279983 | 0.182061 | 1.537856 | 0.124084 | 423.8833 | P | 250 |
| *Johngarthia lagostoma* | Ilha do Meio | lam(landscapeCover) gamma(.) p(totalRainfallThreeMonths+moonPhase) iota(.) | lambda | (Intercept) | 4.306621 | 0.321849 | 13.38087 | 7.82E-41 | 672.0895 | ZIP | 650 |
| | | | lambda | landscapeCoveropen | -0.4134 | 0.212487 | -1.94555 | 0.051709 | 672.0895 | ZIP | 650 |
| | | | lambda | landscapeCovertreed | 1.316558 | 0.182576 | 7.211004 | 5.55E-13 | 672.0895 | ZIP | 650 |
| | | | gamma | (Intercept) | 0.096264 | 0.024073 | 3.998883 | 6.36E-05 | 672.0895 | ZIP | 650 |
| | | | det | (Intercept) | -3.81571 | 0.288732 | -13.2154 | 7.15E-40 | 672.0895 | ZIP | 650 |
| | | | det | scale(totalRainfallThreeMonths) | 0.732966 | 0.060521 | 12.11103 | 9.23E-34 | 672.0895 | ZIP | 650 |
| | | | det | scale(moonPhase) | -0.39391 | 0.069716 | -5.65021 | 1.60E-08 | 672.0895 | ZIP | 650 |
| | | | psi | psi | -2.46835 | 0.75721 | -3.25979 | 0.001115 | 672.0895 | ZIP | 650 |
| | Ilha Rata | lam(.) gamma(.) p(totalRainfallThreeMonths+moonPhase) iota(.) | lambda | (Intercept) | 2.682739 | 0.299123 | 8.968692 | 3.00E-19 | 471.8631 | ZIP | 450 |
| | | | gamma | (Intercept) | 0.241543 | 0.040056 | 6.030154 | 1.64E-09 | 471.8631 | ZIP | 450 |
| | | | det | (Intercept) | -1.85314 | 0.335904 | -5.51689 | 3.45E-08 | 471.8631 | ZIP | 450 |
| | | | det | scale(totalRainfallThreeMonths) | 0.130363 | 0.073027 | 1.78514 | 0.074239 | 471.8631 | ZIP | 450 |
| | | | det | scale(moonPhase) | -0.17314 | 0.068049 | -2.54441 | 0.010946 | 471.8631 | ZIP | 450 |
| | | | psi | psi | -1.32592 | 0.600379 | -2.20847 | 0.027212 | 471.8631 | ZIP | 450 |
| *Sula dactylatra* | Ilha do Meio | lam(.) gamma(TSE) p(.) iota(.) | lambda | (Intercept) | 4.914824 | 0.154351 | 31.84193 | 1.70E-222 | 598.8573 | P | 754 |
| | | | gamma | (Intercept) | 0.231095 | 0.024625 | 9.384399 | 6.33E-21 | 598.8573 | P | 754 |
| | | | gamma | scale(TSE) | 0.143032 | 0.038033 | 3.760787 | 0.000169 | 598.8573 | P | 754 |
| | | | det | (Intercept) | -0.79571 | 0.22122 | -3.59691 | 0.000322 | 598.8573 | P | 754 |
| | Ilha Rata | lam(.) gamma(.) p(.) iota(.) | lambda | (Intercept) | 6.764202 | 0.116119 | 58.25242 | 0 | 147.4354 | NB | 2319 |
| | | | gamma | (Intercept) | 0.318583 | 0.035816 | 8.894981 | 5.84E-19 | 147.4354 | NB | 2319 |
| | | | det | (Intercept) | -1.98912 | 0.089925 | -22.1197 | 2.04E-108 | 147.4354 | NB | 2319 |
| | | | alpha | alpha | 12.09775 | 32.56222 | 0.371527 | 0.710245 | 147.4354 | NB | 2319 |

**Table C3.** Population density through time, detection probabilities (i.e., observation), and population growth (i.e., gamma or the finite rate of increase, normally referred to as lambda in ecology) derived from fixed-radius point counts on Ilha do Meio and Ilha Rata (Brazil) for native species. Black rats (*Rattus rattus*) were controlled and eventually eradicated on Ilha do Meio (between cycles 3 and 4), while Ilha Rata, where black rat populations exist and are stable, was used to control indices for seasonality and other regional factors. During cycle 3, rat capture-recapture and native species' population census were not performed and are therefore not shown. Data collection on *E. ridleyana* was maintained for further cycles, every May and October in 2019, and 2021 to 2023. In some cycles, wildlife monitoring was performed twice or three times, resulting in decimal numbers in the cycles' column. Estimates were averaged across site types (i.e., when different site covers presented different estimates) and values in parenthesis represent confidence interval at 95%.

| *Species* | *Island* | *Cycle* | *Detection probability* | *Initial Population Density (m2)* | *Population Growth* |
|---|---|---|---|---|---|
| *Trachylepis atlantica* | Ilha do Meio | 1 | 0.048 (0.005 - 0.285) | 3.759 (0.106 - 10.858) | 0.568 (0.438 - 0.737) |
| | | 2 | 0.053 (0.015 - 0.285) | 2.136 (0.046 - 8.002) | 0.937 (0.845 - 1.04) |
| | | 2.1 | 0.053 (0.015 - 0.285) | 2.002 (0.039 - 8.322) | 0.944 (0.852 - 1.047) |
| | | 4 | 0.053 (0.015 - 0.285) | 1.891 (0.033 - 8.712) | 2.063 (1.52 - 2.799) |
| | | 5 | 0.051 (0.012 - 0.285) | 3.9 (0.051 - 24.383) | NA |
| | Ilha Rata | 1 | 0.005 (0 - 0.529) | 6.383 (6.383 - 6.383) | 0.993 (0.848 - 1.163) |
| | | 2 | 0.005 (0 - 0.529) | 6.338 (5.411 - 7.423) | 0.993 (0.848 - 1.163) |
| | | 2.1 | 0.011 (0 - 0.705) | 6.293 (4.587 - 8.634) | 0.993 (0.848 - 1.163) |
| | | 4 | 0.011 (0 - 0.705) | 6.249 (3.889 - 10.042) | 0.993 (0.848 - 1.163) |
| | | 5 | 0.011 (0 - 0.722) | 6.205 (3.297 - 11.679) | 0.993 (0.848 - 1.163) |
| | | 5.1 | 0.011 (0 - 0.705) | 6.162 (2.795 - 13.583) | NA |
| *Johngarthia lagostoma* | Ilha do Meio | 1 | 0.074 (0.041 - 0.131) | 1.565 (0.576 - 3.249) | 1.101 (1.05 - 1.154) |
| | | 1.1 | 0.042 (0.024 - 0.074) | 1.723 (0.605 - 3.75) | 1.101 (1.05 - 1.154) |
| | | 2 | 0.01 (0.005 - 0.017) | 1.897 (0.635 - 4.328) | 1.101 (1.05 - 1.154) |
| | | 2.1 | 0.007 (0.004 - 0.013) | 2.089 (0.667 - 4.996) | 1.101 (1.05 - 1.154) |
| | | 4 | 0.017 (0.01 - 0.03) | 2.3 (0.701 - 5.766) | 1.101 (1.05 - 1.154) |
| | | 4.1 | 0.015 (0.009 - 0.027) | 2.533 (0.736 - 6.656) | 1.101 (1.05 - 1.154) |
| | | 5 | 0.037 (0.021 - 0.065) | 2.789 (0.773 - 7.682) | NA |
| | Ilha Rata | 1 | 0.133 (0.069 - 0.241) | 0.147 (0.147 - 0.147) | 1.273 (1.177 - 1.377) |
| | | 2 | 0.144 (0.079 - 0.247) | 0.187 (0.173 - 0.203) | 1.273 (1.177 - 1.377) |
| | | 2.1 | 0.14 (0.077 - 0.241) | 0.239 (0.204 - 0.279) | 1.273 (1.177 - 1.377) |
| | | 2.2 | 0.134 (0.074 - 0.23) | 0.304 (0.24 - 0.384) | 1.273 (1.177 - 1.377) |
| | | 4 | 0.097 (0.052 - 0.172) | 0.387 (0.282 - 0.529) | 1.273 (1.177 - 1.377) |
| | | 5 | 0.176 (0.093 - 0.307) | 0.492 (0.332 - 0.729) | NA |
| *Elaenia ridleyana* | Ilha do Meio | 1 | 0.49 (0.392 - 0.589) | 0.099 (0.099 - 0.099) | 0.689 (0.463 - 1.025) |
| | | 2 | 0.49 (0.392 - 0.589) | 0.068 (0.046 - 0.101) | 0.704 (0.478 - 1.039) |
| | | 4 | 0.49 (0.392 - 0.589) | 0.048 (0.022 - 0.105) | 1.183 (0.881 - 1.587) |
| | | 5 | 0.49 (0.392 - 0.589) | 0.057 (0.019 - 0.167) | 1.299 (0.949 - 1.779) |
| | | 6 | 0.49 (0.392 - 0.589) | 0.074 (0.018 - 0.297) | 1.457 (1.04 - 2.039) |

*Table C3 cont.*

| | | | | | |
|---|---|---|---|---|---|
| | | 7 | 0.49 (0.392 - 0.589) | 0.107 (0.019 - 0.605) | 1.457 (1.041 - 2.04) |
| | | 8 | 0.49 (0.392 - 0.589) | 0.156 (0.02 - 1.233) | 0.799 (0.586 - 1.089) |
| | | 9 | 0.49 (0.392 - 0.589) | 0.125 (0.012 - 1.344) | 0.795 (0.582 - 1.086) |
| | | 10 | 0.49 (0.392 - 0.589) | 0.099 (0.007 - 1.459) | 0.409 (0.216 - 0.776) |
| | | 11 | 0.49 (0.392 - 0.589) | 0.041 (0.001 - 1.133) | 1.848 (1.156 - 2.956) |
| | | 12 | 0.49 (0.392 - 0.589) | 0.075 (0.002 - 3.348) | 0.709 (0.447 - 1.125) |
| | | 13 | 0.49 (0.392 - 0.589) | 0.053 (0.001 - 3.766) | 1.979 (1.141 - 3.433) |
| | | 14 | 0.49 (0.392 - 0.589) | 0.105 (0.001 - 12.931) | 0.724 (0.425 - 1.232) |
| | | 15 | 0.49 (0.392 - 0.589) | 0.076 (0 - 15.93) | NA |
| | Ilha Rata | 1 | 0.57 (0.481 - 0.654) | 0.062 (0.062 - 0.062) | 0.962 (0.739 - 1.251) |
| | | 2 | 0.57 (0.481 - 0.654) | 0.059 (0.046 - 0.077) | 0.96 (0.738 - 1.249) |
| | | 4 | 0.57 (0.481 - 0.654) | 0.057 (0.034 - 0.097) | 1.408 (0.991 - 2.001) |
| | | 5 | 0.57 (0.481 - 0.654) | 0.08 (0.033 - 0.193) | 1.407 (0.991 - 1.999) |
| | | 6 | 0.57 (0.481 - 0.654) | 0.113 (0.033 - 0.387) | 1.422 (0.995 - 2.034) |
| | | 7 | 0.57 (0.481 - 0.654) | 0.161 (0.033 - 0.787) | 1.416 (0.993 - 2.019) |
| | | 8 | 0.57 (0.481 - 0.654) | 0.228 (0.033 - 1.588) | 0.928 (0.705 - 1.222) |
| | | 9 | 0.57 (0.481 - 0.654) | 0.212 (0.023 - 1.94) | 0.928 (0.705 - 1.221) |
| | | 10 | 0.57 (0.481 - 0.654) | 0.196 (0.016 - 2.37) | 0.565 (0.308 - 1.035) |
| | | 11 | 0.57 (0.481 - 0.654) | 0.111 (0.005 - 2.453) | 1.305 (0.959 - 1.775) |
| | | 12 | 0.57 (0.481 - 0.654) | 0.145 (0.005 - 4.354) | 0.667 (0.414 - 1.074) |
| | | 13 | 0.57 (0.481 - 0.654) | 0.097 (0.002 - 4.677) | 1.282 (0.951 - 1.729) |
| | | 14 | 0.57 (0.481 - 0.654) | 0.124 (0.002 - 8.086) | 0.631 (0.376 - 1.06) |
| | | 15 | 0.57 (0.481 - 0.654) | 0.078 (0.001 - 8.57) | NA |
| *Sula dactylatra* | Ilha do Meio | 1 | 0.311 (0.226 - 0.41) | 0.001 (0.001 - 0.001) | 1.074 (0.968 - 1.19) |
| | | 2 | 0.311 (0.226 - 0.41) | 0.001 (0.001 - 0.001) | 1.235 (1.173 - 1.3) |
| | | 4 | 0.311 (0.226 - 0.41) | 0.001 (0.001 - 0.001) | 1.509 (1.368 - 1.664) |
| | | 5 | 0.311 (0.226 - 0.41) | 0.002 (0.001 - 0.002) | NA |
| | Ilha Rata | 1 | 0.12 (0.103 - 0.14) | 0.005 (0.005 - 0.005) | 1.375 (1.282 - 1.475) |
| | | 2 | 0.12 (0.103 - 0.14) | 0.007 (0.006 - 0.007) | 1.375 (1.282 - 1.475) |
| | | 4 | 0.12 (0.103 - 0.14) | 0.009 (0.008 - 0.01) | 1.375 (1.282 - 1.475) |
| | | 5 | 0.12 (0.103 - 0.14) | 0.013 (0.01 - 0.015) | NA |
