## Appendix D for "Swiftly squeaky clean: lessons learned from eradicating an overpopulation of rats on an island of constraints"

**Table D1.** Results of a literature search conducted using the Scholarcy (https://www.scholarcy.com/) tool with the search string: *((("adaptive") AND ("management")) AND ("invasive")) AND ("species")*. The search aimed to identify studies that applied adaptive management in the context of invasive species. Each entry provides key details about the study, such as title, author, year of publication, whether adaptive management was used, whether the study was conducted on an island, if it was in a tropical setting, and whether it followed the six adaptive management points outlined by Westgate et al. (2013). Additionally, the Digital Object Identifier (DOI) for each study is listed for reference.

| Title | Author | Year of publication | Uses AM? | Done on Island? | Tropical? | Follows Westgate 6 points? | DOI |
|---|---|---|---|---|---|---|---|
| **Successful recovery of North Island kokako Callaeas cinerea wilsoni populations, by adaptive management** | Innes, John | 1999 | Yes | No | No | 1 to 5 | 10.1016/S0006-3207(98)00053-6 |
| **Large scale predator control improves the productivity of a rare New Zealand riverine duck** | Whitehead, Amy L. | 2008 | Yes | No | No | 1 to 5 | 10.1016/j.biocon.2008.08.013 |
| **Energy and water use by invasive goats (Capra hircus) in an Australian rangeland, and a caution against using broad-scale allometry to predict species-specific requirements.** | Munn, A. J. | 2012 | No | | | | 10.1016/j.cbpa.2011.10.027 |
| **A battle lost? Report on two centuries of invasion and management of Lantana camara L. in Australia, India and South Africa.** | Bhagwat, Shonil A. | 2012 | No | | | | 10.1371/journal.pone.0032407 |
| **Mutual interactions between an invasive bark beetle and its associated fungi.** | Wang, B. | 2012 | No | | | | 10.1017/S000748531100037X |
| **Combining structured decision making and value-of-information** | Moore, Joslin L. | 2012 | No | | | | 10.1111/j.1523-1739.2012.01907.x |

| | | | | | | | |
|---|---|---|---|---|---|---|---|
| **analyses to identify robust management strategies.** | | | | | | | |
| **Anthropogenically induced adaptation to invade (AIAI): contemporary adaptation to human-altered habitats within the native range can promote invasions.** | Hufbauer, Ruth A. | 2012 | No | | | | 10.1111/j.1752-4571.2011.00211.x |
| **Climate-Driven Reshuffling of Species and Genes: Potential Conservation Roles for Species Translocations and Recombinant Hybrid Genotypes.** | Scriber, Jon Mark | 2013 | No | | | | 10.3390/insects5010001 |
| **Climate change expands the spatial extent and duration of preferred thermal habitat for lake Superior fishes.** | Cline, Timothy J. | 2013 | No | | | | 10.1371/journal.pone.0062279 |
| **Genomic patterns of introgression in rainbow and westslope cutthroat trout illuminated by overlapping paired-end RAD sequencing.** | Hohenlohe, Paul A. | 2013 | No | | | | 10.1111/mec.12239 |
| **Nutrient limitation of native and invasive N2-fixing plants in northwest prairies.** | Thorpe, Andrea S. | 2013 | No | | | | 10.1371/journal.pone.0084593 |
| **Migration and dispersal may drive to high genetic variation and significant genetic mixing: the case of two agriculturally important, continental hoverflies (Episyrphus balteatus and Sphaerophoria scripta).** | Raymond, Lucie | 2013 | No | | | | 10.1111/mec.12483 |
| **Towards the planning and design of disturbance patterns across scales to counter biological invasions.** | Zurlini, Giovanni | 2013 | No | | | | 10.1016/j.jenvman.2013.05.006 |
| **High Variation in Single Nucleotide Polymorphisms (SNPs) and Insertions/Deletions (Indels) in the Highly Invasive Bemisia tabaci (Gennadius) (Hemiptera:** | Lü, Z. C. | 2013 | No | | | | 10.1007/s13744-013-0152-2 |

| | | | | | | | |
|---|---|---|---|---|---|---|---|
| **Aleyrodidae) Middle East-Asia Minor 1 (MEAM1).** | | | | | | | |
| **Novel microsatellite DNA markers indicate strict parthenogenesis and few genotypes in the invasive willow sawfly Nematus oligospilus.** | Caron, V. | 2013 | No | | | | 10.1017/S0007485312000429 |
| **Stress for invasion success? Temperature stress of preceding generations modifies the response to insecticide stress in an invasive pest insect.** | Piiroinen, Saija | 2013 | No | | | | 10.1111/eva.12001 |
| **Marsh thistle in New York: early detection and rapid response to a recent invader.** | Hinchey, Eamonn | 2013 | No | | | | 10.1111/nyas.12176 |
| **The role of genomics in conservation and reproductive sciences.** | Johnson, Warren E. | 2014 | No | | | | 10.1007/978-1-4939-0820-2_5 |
| **Increased survival and prolonged longevity mainly contribute to the temperature-adaptive evolutionary strategy in invasive Bemisia tabaci (Hemiptera: Aleyrodidae) Middle East Asia Minor 1.** | Lü, Zhi-Chuang | 2014 | No | | | | 10.1093/jisesa/ieu005 |
| **Spinosad and the tomato borer Tuta absoluta: a bioinsecticide, an invasive pest threat, and high insecticide resistance.** | Campos, Mateus R. | 2014 | No | | | | 10.1371/journal.pone.0103235 |
| **Learning from conservation planning for the U.S. National Wildlife Refuges.** | Meretsky, Vicky J. | 2014 | No | | | | 10.1111/cobi.12292 |
| **Phenology research for natural resource management in the United States.** | Enquist, Carolyn A. F. | 2014 | No | | | | 10.1007/s00484-013-0772-6 |
| **Adaptive responses reveal contemporary and future ecotypes in a desert shrub.** | Richardson, Bryce A. | 2014 | No | | | | 10.1890/13-0587.1 |
| **A mid-term analysis of progress toward international biodiversity targets.** | Tittensor, Derek P. | 2014 | No | | | | 10.1126/science.1257484 |

| | | | | | | | | |
|---|---|---|---|---|---|---|---|---|
| **Priority threat management of invasive animals to protect biodiversity under climate change.** | Firn, Jennifer | 2015 | No | | | | | 10.1111/gcb.13034 |
| **Immune Reconstitution After Allogeneic Hematopoietic Stem Cell Transplantation and Association With Occurrence and Outcome of Invasive Aspergillosis.** | Stuehler, Claudia | 2015 | No | | | | | 10.1093/infdis/jiv143 |
| **The potential for biodiversity offsetting to fund effective invasive species control.** | Norton, David A. | 2015 | No | | | | | 10.1111/cobi.12345 |
| **Non-native megaherbivores: the case for novel function to manage plant invasions on islands.** | Hansen, Dennis M. | 2015 | No | | | | | 10.1093/aobpla/plv085 |
| **Intraspecific Niche Variation Drives Abundance-Occupancy Relationships in Freshwater Fish Communities.** | Faulks, Leanne | 2015 | No | | | | | 10.1086/682004 |
| **Consequences of seed origin and biological invasion for early establishment in restoration of a North American grass species.** | Herget, Mollie E. | 2015 | No | | | | | 10.1371/journal.pone.0119889 |
| **The whole genome sequence of the Mediterranean fruit fly, Ceratitis capitata (Wiedemann), reveals insights into the biology and adaptive evolution of a highly invasive pest species.** | Papanicolaou, Alexie | 2016 | No | | | | | 10.1186/s13059-016-1049-2 |
| **Adaptive management of invasive pests in natural protected areas: the case of Matsucoccus feytaudi in Central Italy.** | Sciarretta, A. | 2016 | Yes | No | No | Yes, but not all were specifically highlighted (i.e., point 2 is not specified -- management options). Note: eradication failed. | | 10.1017/S0007485315000851 |
| **Biological invasions, ecological resilience and adaptive governance.** | Chaffin, Brian C. | 2016 | No | | | | | 10.1016/j.jenvman.2016.04.040 |

| | | | | | | | |
|---|---|---|---|---|---|---|---|
| **Targeted gene flow for conservation.** | Kelly, Ella | 2016 | No | | | | 10.1111/cobi.12623 |
| **Diagnostic of Fungal Infections Related to Biofilms.** | Sanguinetti, Maurizio | 2016 | No | | | | 10.1007/5584_2016_9 |
| **Improving credibility and transparency of conservation impact evaluations through the partial identification approach.** | McConnachie, Matthew M. | 2016 | No | | | | 10.1111/cobi.12610 |
| **Evidence of Subdivisions on Evolutionary Timescales in a Large, Declining Marsupial Distributed across a Phylogeographic Barrier.** | Alpers, Deryn L. | 2016 | No | | | | 10.1371/journal.pone.0162789 |
| **Controlling range expansion in habitat networks by adaptively targeting source populations.** | Hock, Karlo | 2016 | No | | | | 10.1111/cobi.12665 |
| **Traditional Mapuche ecological knowledge in Patagonia, Argentina: fishes and other living beings inhabiting continental waters, as a reflection of processes of change.** | Aigo, Juana | 2016 | No | | | | 10.1186/s13002-016-0130-y |
| **Enhancing the effectiveness of biological control programs of invasive species through a more comprehensive pest management approach.** | DiTomaso, Joseph M. | 2017 | No | | | | 10.1002/ps.4347 |
| **Erratum to: The whole genome sequence of the Mediterranean fruit fly, Ceratitis capitata (Wiedemann), reveals insights into the biology and adaptive evolution of a highly invasive pest species.** | Papanicolaou, Alexie | 2017 | No | | | | 10.1186/s13059-017-1155-9 |
| **A quick and robust MHC typing method for free-ranging and captive primate species.** | de Groot, N. | 2017 | No | | | | 10.1007/s00251-016-0968-0 |
| **Selection of invasive wild pig countermeasures using multicriteria decision analysis.** | Brondum, Matthew C. | 2017 | No | | | | 10.1016/j.scitotenv.2016.09.155 |

| | | | | | | | | |
|---|---|---|---|---|---|---|---|---|
| **Eradicating the grey squirrel Sciurus carolinensis from urban areas: an innovative decision-making approach based on lessons learnt in Italy.** | La Morgia, Valentina | 2017 | No | | | | | 10.1002/ps.4352 |
| **GMDPtoolbox: A Matlab library for designing spatial management policies. Application to the long-term collective management of an airborne disease.** | Cros, Marie-Josée | 2017 | No | | | | | 10.1371/journal.pone.0186014 |
| **Using Landscape Genetics Simulations for Planting Blister Rust Resistant Whitebark Pine in the US Northern Rocky Mountains.** | Landguth, Erin L. | 2017 | No | | | | | 10.3389/fgene.2017.00009 |
| **How to improve threatened species management: An Australian perspective.** | Scheele, B. C. | 2018 | No | | | | | 10.1016/j.jenvman.2018.06.084 |
| **Resilience and Adaptation: Yukon River Watershed Contaminant Risk Indicators.** | Duffy, Lawrence | 2018 | No | | | | | 10.1155/2018/8421513 |
| **High-Throughput Analysis Reveals Seasonal Variation of the Gut Microbiota Composition Within Forest Musk Deer (Moschus berezovskii).** | Hu, Xiaolong | 2018 | No | | | | | 10.3389/fmicb.2018.01674 |
| **Modelling the invasion history of Sinanodonta woodiana in Europe: Tracking the routes of a sedentary aquatic invader with mobile parasitic larvae.** | Konečný, Adam | 2018 | No | | | | | 10.1111/eva.12700 |
| **The role of invasive alien species in shaping local livelihoods and human well-being: A review.** | Shackleton, Ross T. | 2019 | No | | | | | 10.1016/j.jenvman.2018.05.007 |
| **Recent research status of Bactrocera dorsalis: Insights from resistance mechanisms and population structure.** | Wei, Dan-Dan | 2019 | No | | | | | 10.1002/arch.21601 |
| **Parallel introgression and selection on introduced alleles in a native species.** | Bay, Rachael A. | 2019 | No | | | | | 10.1111/mec.15097 |

| | | | | | | | |
|---|---|---|---|---|---|---|---|
| **Stress in captive Blue-fronted parrots (Amazona aestiva): the animalists' tale.** | Vidal, Alan Chesna | 2019 | No | | | | 10.1093/conphys/coz097 |
| **A Novel Cold-Adaptive Endo-1,4-β-Glucanase From Burkholderia pyrrocinia JK-SH007: Gene Expression and Characterization of the Enzyme and Mode of Action.** | Chen, Feifei | 2019 | No | | | | 10.3389/fmicb.2019.03137 |
| **Targeted gene flow and rapid adaptation in an endangered marsupial.** | Kelly, Ella | 2019 | No | | | | 10.1111/cobi.13149 |
| **Enhancement of oxidative stress contributes to increased pathogenicity of the invasive pine wood nematode.** | Zhang, Wei | 2019 | No | | | | 10.1098/rstb.2018.0323 |
| **Invasion origin, rapid population expansion, and the lack of genetic structure of cotton bollworm (Helicoverpa armigera) in the Americas.** | Gonçalves, Rogério Martins | 2019 | No | | | | 10.1002/ece3.5123 |
| **Rapid growth and defence evolution following multiple introductions.** | van Boheemen, Lotte A. | 2019 | No | | | | 10.1002/ece3.5275 |
| **Risk-Based and Adaptive Invasive Mosquito Surveillance at Lucky Bamboo and Used Tire Importers in the Netherlands.** | Ibáñez-Justicia, Adolfo | 2020 | No | | | | 10.2987/20-6914.1 |
| **Integrated Methods for Monitoring the Invasive Potential and Management of Heracleum mantegazzianum (giant hogweed) in Switzerland.** | Shackleton, Ross T. | 2020 | No | | | | 10.1007/s00267-020-01282-9 |
| **Testing the adaptive advantage of a threatened species over an invasive species using a stochastic population model.** | Brown, Timothy R. | 2020 | No | | | | 10.1016/j.jenvman.2020.110524 |
| **Our Wild Companions: Domestic cats in the Anthropocene.** | Crowley, Sarah L. | 2020 | No | | | | 10.1016/j.tree.2020.01.008 |
| **Abundance of invasive grasses is dependent on fire regime and** | Damasceno, Gabriella | 2020 | No | | | | 10.1016/j.jenvman.2020.111016 |

| | | | | | | | |
|---|---|---|---|---|---|---|---|
| **climatic conditions in tropical savannas.** | | | | | | | |
| **Forest genomics: Advancing climate adaptation, forest health, productivity, and conservation.** | Isabel, Nathalie | 2020 | No | | | | 10.1111/eva.12902 |
| **Reticulate evolution as a management challenge: Patterns of admixture with phylogenetic distance in endemic fishes of western North America.** | Bangs, Max R. | 2020 | No | | | | 10.1111/eva.13042 |
| **Hitchhikers on floats to Arctic freshwater: Private aviation and recreation loss from aquatic invasion.** | Schwoerer, Tobias | 2020 | No | | | | 10.1007/s13280-019-01295-7 |
| **Origin of resistance to pyrethroids in the redlegged earth mite (Halotydeus destructor) in Australia: repeated local evolution and migration.** | Yang, Qiong | 2020 | No | | | | 10.1002/ps.5538 |
| **Dramatic long-term restoration of an oak woodland due to multiple, sustained management treatments.** | Glennemeier, Karen | 2020 | No | | | | 10.1371/journal.pone.0241061 |
| **Spatial optimization of invasive species control informed by management practices.** | Nishimoto, Makoto | 2021 | No | | | | 10.1002/eap.2261 |
| **Kentucky Bluegrass Invasion in the Northern Great Plains and Prospective Management Approaches to Mitigate Its Spread.** | Palit, Rakhi | 2021 | No | | | | 10.3390/plants10040817 |
| **Fishery reforms for the management of non-indigenous species.** | Kleitou, Periklis | 2021 | No | | | | 10.1016/j.jenvman.2020.111690 |
| **What Are the Keys to the Adaptive Success of European Wild Rabbit (Oryctolagus cuniculus) in the Iberian Peninsula?** | Marín-García, Pablo Jesús | 2021 | No | | | | 10.3390/ani11082453 |
| **Adaptive population structure shifts in invasive parasitic mites, Varroa destructor.** | Moro, Arrigo | 2021 | No | | | | 10.1002/ece3.7272 |
| **Rapid genetic adaptation to recently colonized environments is** | Yin, Xiaoshen | 2021 | No | | | | 10.1186/s12864-021-07553-x |

| | | | | | | | |
|---|---|---|---|---|---|---|---|
| **driven by genes underlying life history traits.** | | | | | | | |
| **The Impact of Climate Change on Forest Development: A Sustainable Approach to Management Models Applied to Mediterranean-Type Climate Regions.** | Nunes, Leonel J. R. | 2021 | No | | | | 10.3390/plants11010069 |
| **Raiders of the last ark: the impacts of feral cats on small mammals in Tasmanian forest ecosystems.** | Lazenby, B. T. | 2021 | No | | | | 10.1002/eap.2362 |
| **Genomic analysis unveils mechanisms of northward invasion and signatures of plateau adaptation in the Asian house rat.** | Chen, Yi | 2021 | No | | | | 10.1111/mec.16194 |
| **Evolutionary genomics of endangered Hawaiian tree snails (Achatinellidae: Achatinellinae) for conservation of adaptive capacity.** | Price, Melissa R. | 2021 | No | | | | 10.7717/peerj.10993 |
| **Prolonged Treatment with Grains of Paradise (Aframomum melegueta) Extract Recruits Adaptive Thermogenesis and Reduces Body Fat in Humans with Low Brown Fat Activity.** | Yoneshiro, Takeshi | 2021 | No |  |  |  | 10.3177/jnsv.67.99 |
| **Population genetic structure of raccoons as a consequence of multiple introductions and range expansion in the Boso Peninsula, Japan.** | Hirose, Miki | 2021 | No | | | | 10.1038/s41598-021-98029-1 |
| **Risk management recommendations for environmental releases of gene drive modified insects.** | Devos, Yann | 2022 | No | | | | 10.1016/j.biotechadv.2021.107807 |
| **The Prediction of Distribution of the Invasive Fallopia Taxa in Slovakia.** | Gašparovičová, Petra | 2022 | No | | | | 10.3390/plants11111484 |
| **Vegetation, water infiltration, and soil carbon response to Adaptive Multi-Paddock and Conventional grazing in Southeastern USA ranches.** | Apfelbaum, Steven I. | 2022 | No | | | | 10.1016/j.jenvman.2022.114576 |

| | | | | | | | |
|---|---|---|---|---|---|---|---|
| **Effectiveness of front line and emerging fungal disease prevention and control interventions and opportunities to address appropriate eco-sustainable solutions.** | Garvey, Mary | 2022 | No | | | | 10.1016/j.scitotenv.2022.158284 |
| **Effect of microfibers combined with UV-B and drought on plant community.** | Deng, Jiaojiao | 2022 | No | | | | 10.1016/j.chemosphere.2021.132413 |
| **(Epi)genomic adaptation driven by fine geographical scale environmental heterogeneity after recent biological invasions.** | Chen, Yiyong | 2022 | No | | | | 10.1002/eap.2772 |
| **Applying early warning indicators to predict critical transitions in a lake undergoing multiple changes.** | Rohde, Elizabeth | 2022 | No | | | | 10.1002/eap.2685 |
| **Habitat suitability and connectivity modeling predict genetic population structure and priority control areas for invasive nutria (Myocastor coypus) in a temperate river basin.** | Kang, Wanmo | 2022 | No | | | | 10.1371/journal.pone.0279082 |
| **Historical museum samples enable the examination of divergent and parallel evolution during invasion.** | Stuart, Katarina C. | 2022 | No | | | | 10.1111/mec.16353 |
| **Impact of roadside burning on genetic diversity in a high-biomass invasive grass.** | Di, Binyin | 2022 | No | | | | 10.1111/eva.13369 |
| **Individual and population-scale carbon and nitrogen isotopic values of Procambarusclarkii in invaded freshwater ecosystems.** | Di Muri, Cristina | 2022 | No | | | | 10.3897/BDJ.10.e94411 |
| **Native species exhibit physiological habituation to invaders: a reason for hope.** | Santicchia, Francesca | 2022 | No | | | | 10.1098/rspb.2022.1022 |
| **Phenotypes and environment predict seedling survival for seven co-occurring Great Basin plant taxa growing with invasive grass.** | Agneray, Alison C. | 2022 | No | | | | 10.1002/ece3.8870 |
| **Adaptive risk-based targeted surveillance for foreign animal** | Miller, Ryan S. | 2022 | No | | | | 10.1111/tbed.14576 |

| | | | | | | | |
|---|---|---|---|---|---|---|---|
| **diseases at the wildlife-livestock interface.** | | | | | | | |
| **Remodeling the bladder tumor immune microenvironment by mycobacterial species with changes in their cell envelope composition.** | Senserrich, Jordi | 2022 | No | | | | 10.3389/fimmu.2022.993401 |
| **Resistance Bioassays and Allele Characterization Inform Analysis of Spodoptera frugiperda (Lepidoptera: Noctuidae) Introduction Pathways in Asia and Australia.** | Tay, W. T. | 2022 | No | | | | 10.1093/jee/toac151 |
| **The Role of Intestinal Microbial Metabolites in the Immunity of Equine Animals Infected With Horse Botflies.** | Hu, Dini | 2022 | No | | | | 10.3389/fvets.2022.832062 |
| **Chromosome-level genome assembly and population genomic analyses provide insights into adaptive evolution of the red turpentine beetle, Dendroctonus valens.** | Liu, Zhudong | 2022 | No | | | | 10.1186/s12915-022-01388-y |
| **Are really Nature-Based Solutions sustainable solutions to design future cities in a context of global change? Discussion about the vulnerability of these new solutions and their probable unsustainable implementation.** | Duffaut, Chloé | 2022 | No | | | | 10.1016/j.scitotenv.2022.158535 |
| **Evaluation of thermoregulation of different pine organs in early spring and estimation of heat reward for the western conifer seed bug (Leptoglossus occidentalis) on male cones.** | Kitajima, Ryotaro | 2022 | No | | | | 10.1371/journal.pone.0272565 |
| **The high invasion success of fall armyworm is related to life-history strategies across a range of stressful temperatures.** | Wu, Pengxiang | 2022 | No | | | | 10.1002/ps.6867 |
| **COI Haplotyping and Comparative Microbiomics of the** | Awad, Mona | 2022 | No | | | | 10.3390/biology12010027 |

| | | | | | | | |
|---|---|---|---|---|---|---|---|
| **Peach Fruit Fly, an Emerging Pest of Egyptian Olive Orchards.** | | | | | | | |
| **Adaptive ecological knowledge among the Ndjuka Maroons of French Guiana; a case study of two 'invasive species': Melaleuca quinquenervia and Acacia mangium.** | Theys, Johanna | 2023 | No | | | | 10.1186/s13002-023-00602-7 |
| **Adaptive evolution to the natural and anthropogenic environment in a global invasive crop pest, the cotton bollworm.** | Jin, Minghui | 2023 | No | | | | 10.1016/j.xinn.2023.100454 |
| **Disgust in animals and the application of disease avoidance to wildlife management and conservation.** | Sarabian, Cécile | 2023 | No | | | | 10.1111/1365-2656.13903 |
| **Open water dreissenid mussel control projects: lessons learned from a retrospective analysis.** | Dahlberg, Angelique D. | 2023 | No | | | | 10.1038/s41598-023-36522-5 |
| **Exploring vulnerabilities of inland fisheries in Indian context with special reference to climate change and their mitigation and adaptation: a review.** | Paul, Thankam Theresa | 2023 | No | | | | 10.1007/s00484-022-02417-9 |
| **Preadapted to adapt: underpinnings of adaptive plasticity revealed by the downy brome genome.** | Revolinski, Samuel R. | 2023 | No | | | | 10.1038/s42003-023-04620-9 |
| **Gene family expansion analysis and identification of the histone family in Spodoptera frugiperda.** | Gao, Han | 2023 | No | | | | 10.1016/j.cbd.2023.101142 |
| **Rapidly evolved traits enable new conservation tools: perspectives from the cane toad invasion of Australia.** | Shine, Richard | 2023 | No | | | | 10.1093/evolut/qpad102 |
| **Intraspecific trait plasticity to N and P of the wetland invader, Alternanthera philoxeroides under flooded conditions.** | Harms, Nathan E. | 2023 | No | | | | 10.1002/ece3.9966 |

| | | | | | | | |
|---|---|---|---|---|---|---|---|
| **The Invasive Caucasian Populations of the Brown Marmorated Stink Bug Halyomorpha halys (Hemiptera: Heteroptera: Pentatomidae) Rapidly Adapt Their Ecophysiological Traits to the Local Environmental Conditions.** | Reznik, Sergey Ya | 2023 | No | | | | 10.3390/insects14050424 |
| **Incorporating adaptive genomic variation into predictive models for invasion risk assessment.** | Chen, Yiyong | 2024 | No | | | | 10.1016/j.ese.2023.100299 |
| **Adaptive Management in EBIPM: A Key to Success in Invasive Plant Management** | Leffler | 2012 | No | | | | 10.2111/RANGELANDS-D-12-00053.1 |
| **Applied evolutionary biology could aid management of invaded ecosystems** | Oduor | 2015 | No | | | | 10.1890/EHS14-0026.1 |
| **Adaptive management improves decisions about where to search for invasive species** | Rout | 2017 | No | | | | 10.1016/j.biocon.2017.04.009 |
| **Adaptive invasive species distribution models: a framework for modeling incipient invasions** | Uden | 2015 | No | | | | 10.1007/s10530-015-0914-3 |
